# Identification of Hsp90 inhibitors as potential drugs for the treatment of TSC1/TSC2 deficient cancer

**DOI:** 10.1101/2021.02.26.433022

**Authors:** Evelyn M. Mrozek, Vineeta Bajaj, Yanan Guo, Izabela A. Malinowska, Jianming Zhang, David J. Kwiatkowski

**Affiliations:** Cancer Genetics Lab, Pulmonary Medicine Division, Department of Medicine, Brigham and Women’s Hospital and Harvard Medical School, Boston, Massachusetts, United States of America; Department of Cancer Biology, Dana-Farber Cancer Institute, Department of Biological Chemistry and Molecular Pharmacology, Harvard Medical School, Boston, Massachusetts, United States of America

## Abstract

Inactivating mutations in either *TSC1* or *TSC2* cause Tuberous Sclerosis Complex, an autosomal dominant disorder, characterized by multi-system tumor and hamartoma development. Mutation and loss of function of *TSC1* and/or *TSC2* also occur in a variety of sporadic cancers, and rapamycin and related drugs show highly variable treatment benefit in patients with such cancers. The TSC1 and TSC2 proteins function in a complex that inhibits mTORC1, a key regulator of cell growth, which acts to enhance anabolic biosynthetic pathways. In this study, we identified and validated five cancer cell lines with *TSC1* or *TSC2* mutations and performed a kinase inhibitor drug screen with 197 compounds. The five cell lines were sensitive to several mTOR inhibitors, and cell cycle kinase and HSP90 kinase inhibitors. The IC50 for Torin1 and INK128, both mTOR kinase inhibitors, was significantly increased in three TSC2 null cell lines in which TSC2 expression was restored. Rapamycin was significantly more effective than either INK128 or ganetespib (an HSP90 inhibitor) in reducing the growth of TSC2 null SNU-398 cells in a xenograft model. Combination ganetespib-rapamycin showed no significant enhancement of growth suppression over rapamycin. Hence, although HSP90 inhibitors show strong inhibition of TSC1/TSC2 null cell line growth in vitro, ganetespib showed little benefit at standard dosage in vivo. In contrast, rapamycin which showed very modest growth inhibition in vitro was the best agent for in vivo treatment, but did not cause tumor regression, only growth delay.

## Introduction

Tuberous sclerosis complex (TSC) is an autosomal dominant neurocutaneous disorder, which is caused by inactivating mutation either in *TSC1* or *TSC2*. Mutations in either gene cause the same phenotype, although mutations in *TSC1* are associated with milder clinical severity in multiple respects (1, 2). There are multiple highly specific clinical features of TSC including cortical tubers, subependymal nodules, cardiac rhabdomyoma, kidney angiomyolipoma, pulmonary lymphangioleiomyomatosis, facial and ungual angiofibromas (1-4). Although tumors in TSC are histologically benign, they cause life-threatening issues in 10-15% of patients if left untreated (1, 4).

Inactivating *TSC1* and *TSC2* mutations also occur rarely in multiple cancer types. Cancers with higher rates of *TSC1*/*TSC2* mutation include: urothelial carcinoma of the bladder and upper tract, with 6-10% incidence of *TSC1* mutations (5) and perivascular epithelioid cell tumors (PEComa) with up to 50% frequency of *TSC2* and *TSC1* mutations (6). The mechanistic target of rapamycin (mTOR) is a large (2,549 amino acid) protein kinase that occurs in cells in either of two complexes, mTOR complex 1 (mTORC1) and mTORC2, that have overlapping as well as distinct components. They have different roles, and mTORC1 regulates cell growth in part by enhancing anabolic biosynthetic pathways (7, 8). *TSC1* encodes TSC1/hamartin, *TSC2* encodes TSC2/tuberin, and with TBC1D7 the three proteins form the TSC protein complex (9). This TSC protein complex functions to enhance conversion of Rheb-GTP to Rheb-GDP, through the GAP domain of TSC2, which serves to inactivate the mTORC1 kinase (7, 8). Loss of either TSC1 or TSC2 inactivates the TSC protein complex, leading to constitutively active mTORC1 (10). mTORC1 phosphorylates the translational regulators S6 kinases (S6K1 and S6K2) and eukaryotic translation initiation factor 4E binding protein 1 (4E-BP1), as well as many other downstream proteins (11, 12). Both S6K activation and inactivation of 4E-BP1 by phosphorylation are important downstream effectors of mTORC1 activation (7, 8, 13).

Rapamycin, also called Sirolimus, has antiproliferative and immunosuppressive activities. Rapamycin binds to FK-506-binding protein (FKBP12) with high affinity, and rapamycin-FKBP12 binds to mTORC1 to inhibit its kinase activity in an allosteric manner (14). Rapamycin treatment has highly variable effects on mTORC1 kinase activity, as it completely inhibits phosphorylation of S6K, while having relatively little effect on mTORC1 phosphorylation of 4E-BP1 (11). Rapamycin-FKBP12 does not bind to mTORC2 or affect its function directly (15).

Clinically, rapamycin is FDA-approved for both prevention of allograft rejection, and for treatment of lymphangioleiomyomatosis, Drugs closely related to rapamycin are termed rapalogs, and include temsirolimus, everolimus, and deforolimus. Rapalogs have very similar if not identical activity in vivo (16).

Heat shock protein 90 (HSP90) is an ATP-dependent molecular chaperone, which is highly expressed and helps to maintain proteostasis. HSP90 regulates the proper conformation, function and activity of multiple proteins (about 200 ‘client’ proteins) by protecting them from proteasome-mediated degradation. HSP90 expression is upregulated in many forms of cancer, and is thought to promote/enable malignant transformation, tumor progression, invasion, metastasis, and/or angiogenesis (17, 18).

HSP90 inhibition results in proteasome-mediated degradation of protein substrates (19-22). Luminespib (NVP-AUY922) and ganetespib are HSP90 inhibitors, which have been studied in human cancer clinical trials, but are not FDA-approved. They are known to have anti-proliferative, anti-invasive, and pro-apoptotic effects in glioblastomas (23). Ganetespib is reported to have a good safety profile, with adverse effects like fatigue, diarrhea, nausea, vomiting elevated amylase levels, asthenia, anorexia, and hypersensitivity reactions (24), but no liver, ocular, or cardiac toxicities like previous HSP90 inhibitors. So far, no HSP90 inhibitor has been approved for cancer therapy (21).

## Materials and methods

### Cell lines and cell culture

All cell lines were obtained from the Broad Institute of Harvard and MIT. PEER (T-cell acute lymphoblastic leukemia), SNU-878 (hepatocellular carcinoma), SNU-886 (hepatocellular carcinoma), CW-2 (large intestine adenocarcinoma), 23132/87 (stomach adenocarcinoma), MEF-319 (endometrium adenosquamous carcinoma), KM12 (large intestine adenocarcinoma), HEC-151 (endometrium adenocarcinoma), DV-90 (lung adenocarcinoma), OVK18 (ovarian endometrioid carcinoma) and HCC-95 (lung squamous cell carcinoma) were cultured in RPMI 1650 with 10% fetal bovine serum (FBS); CAL-72 (osteosarcoma) and NCI-H1651 (lung adenocarcinoma) in DMEM/F-12 with 10% FBS; MGH-U1 (urinary bladder carcinoma) and HEK-293 (embryonic kidney cells) in DMEM with 10% FBS; and SNU-398 (hepatocellular carcinoma) in RPMI 1650 GlutaMAX medium with 10% FBS. All media was supplemented with 1% penicillin-streptomycin-amphotericin B (Corning). All cell culture was done in a 37 °C humidified incubator in 5% CO_2_. For serum starvation, cells were incubated with standard medium without FBS for 24 hours. For serum stimulation, FBS was added back with a final concentration of 10% for 30 minutes.

### DNA isolation and sequencing

For DNA purification Gentra Puregene Tissue Kit was used (Qiagen). Oligonucleotide primers for sequencing mutated regions of *TSC1* and *TSC2* on the cell lines, where mutations were reported, were designed with Primer3 (25). FastStart PCR Kit (Sigma-Aldrich) was used for PCR, and products were purified using AMPure XP (Beckman Coulter) beads. PCR products were sequenced by Sanger methodology at the High Throughput Sequencing Service, Brigham and Women’s Hospital.

### Multiplex Ligation Dependent Probe Amplification (MLPA) assays

MLPA was performed by standard methods (MRC-Holland, Amsterdam, The Netherlands) using a probe set that covered five tumor suppressor genes: *LKB1, PTEN, CDKN2A, TSC1,* and *TSC2*. 57 of 425 cancer cell lines showed some degree of reduction in signal for one more *TSC1* or *TSC2* probes, and were subject repeat analysis using probe sets specific for either *TSC1* or *TSC2,* the SALSA MLPA probemix P124 TSC1, and the SALSA MLPA probemix P046-D1 TSC2, respectively. Amplification products were run on an ABI 3100Genetic Analyzer (ABI, USA) and electrophoregrams were generated. Peak heights were loaded into a standard Excel file to determine copy number for each probe and sample pair.

### Immunoblotting

Cells were lysed in cell lysis buffer (Cell Signaling Technology) with added protease and phosphatase inhibitor cocktails (Sigma-Aldrich). The protein concentration was determined using a Bradford assay (Boston BioProducts). For immunoblotting, proteins were loaded in 4–12% gradient NuPAGE bis-tris gels (Life Technologies) or home-made 15% polyacrylamide gels, separated by SDS–PAGE and transferred to PVDF membranes. For detection the following primary antibodies were used: mTOR (2972, Cell Signaling Technology), TSC2, C-20 (sc-893, Santa Cruz Biotechnology), TSC1 (4906S, Cell Signaling Technology), pAKT-Ser473 (4060x, Cell Signaling Technology), AKT1, C-20 (J2810, Santa Cruz Biotechnology), pS6K-Thr389 (9234S, Santa Cruz Biotechnology), S6 kinase, C-18 (D0506, Santa Cruz Biotechnology), pS6-Ser235/236 (4857S, Cell Signaling Technology), pS6-Ser240/244 (2215L, Cell Signaling Technology), GAPDH (GR9686I, Abcam), 4E-BP1 (9452L, Cell Signaling Technology), p4E-BP1-Ser65, (9451S, Cell Signaling Technology), p4E-BP1-Thr37/T46 (2855, Cell Signaling Technology), peIF2α-Ser51 (9721S, Cell Signaling Technology), BiP (3183S, Cell Signaling Technology), cleaved caspase-3 (9664S, Cell Signaling Technology) and β-Actin (4970, Cell Signaling Technology).

Secondary antibodies were anti-mouse, anti-rabbit, and anti-goat (Santa Cruz Biotechnology) conjugated to horseradish peroxidase, and were used at 1:3000 dilution. Immunoreactive bands were detected by chemiluminescence (super signal west pico and femto chemiluminescent, Thermo Fisher Scientific) using the G:Box:iChemiXT imager (Syngene).

### Kinase inhibitor library screen and IC50 determination

The cell lines SNU-878, SNU-886, SNU-398, CAL-72, and PEER were screened for kinase inhibitors using a kinase inhibitor-focused library (LINCS). The LINCS library contained 197 kinase inhibitors, from a diverse ATP competitive kinase inhibitor set. These kinase inhibitors were shown to be relatively potent and selective towards a narrow range of targets. 2000 adherent cells and 3000 suspension cells were plated with 50 µL medium in each well of a 384 microplate. Drugs with a concentration of 660 nM were added the same day. After 48 hours cellular proliferation was determined using CellTiter-Glo (Promega). Drugs identified as having a significant effect on this screen were tested in greater detail. 2000 adherent or 4000 suspension cells were plated on day 0 on a 96 well plate with 100 µL medium, except on the marginal wells. Drugs were added on day 1, which were serially diluted three-fold from 10 μM to 1.5 nM. Cell viability was determined after 72 hours of treatment by adding 20 µL CellTiter-Glo. Cell viability was determined using XLfit4.0 software. IC_50_ values were calculated using Graphpad Prism as the drug concentration that reduced cell viability by 50% compared to untreated cells. Rapamycin was purchased from LC laboratories and ganetespib from Synta Pharmaceuticals.

### TSC2 add-back

To add back TSC2 to cells lacking a functional TSC2 expression (SNU-878, SNU-886, and SNU-398 cells), pMSCV and pMSCV-TSC2 (containing the TSC2 cDNA) plasmids were used. HEK293T cells were transfected with pCL Ampho retrovirus packaging vector (Imgenex) and pMSCV-EV (empty vector) or pMSCV-TSC2, with selection to generate retrovirus. Retrovirus was then added to each of the SNU cell lines. Neomycin was used as a selection antibiotic.

### Mouse xenograft studies

All procedures were carried out according to the Guide for the Humane Use and Care of Laboratory Animals, and the experiments were approved by the Animal Care and Use Committee of the Children’s Hospital, Boston, MA, USA (protocol reference number: 1589).

Immunodeficient strain CB17SC-M scid (C.B-*Igh-1^b^*/IcrTac-*Prkdc^scid^*) mice were used to generate xenograft tumors. Mice were purchased at the age of 3-4 weeks from Taconic, Germantown, NY, USA. Mice were 6 to 8 weeks old and their weight was between 17 to 25 g when 3 x 10^6^ tumor cells of the cancer cell line SNU-398 in 100 µL medium mixed with 100 µL matrigel were injected subcutaneously in both back flank regions.

Xenograft tumor nodules were followed 3x weekly, and when they reached a minimum diameter of 4mm (14-30 days after injection), drug treatments were initiated. Mice were treated either with INK128 (Intellikine Inc.), rapamycin (LC laboratories), or ganetespib (Synta Pharmaceuticals); or vehicle; or a combination of ganetespib and INK128, or ganetespib and rapamycin. Bodyweight was measured at least 3 times a week. Tumor size was measured with an electronic digital caliper in 2 dimensions (volume= a^2^xb/2, with a being the greater diameter). For drug administration, INK128 powder was dissolved daily in vehicle (5% NMP, 15% polyvinylpyrrolidone K30, and 80% water); and was administered by gavage 5 days/week at a dose of 1mg/kg and a volume of 100 µL. Rapamycin was prepared as a 20 mg/mL stock solution using 100% ethanol, and was mixed daily with sterile vehicle (0.25% PEG-200, 0.25% Tween-80, and water to achieve a volume of 200 µL for injection. Rapamycin was administered by intraperitoneal injection 3 times/week in a dose of 3 mg/kg. Ganetespib powder was dissolved in DMSO to concentration 50 mg/mL, with heating to 55 °C. Ganetespib stock solution was diluted 1:10 in 20% cremophor RH40 (CrRH40)/80% dextrose (D5W). Ganetespib was administered by tail vein injection once/week in a dose of 50 mg/kg. Control mice received vehicle treatment for one of these three drugs.

### Immunohistochemistry (IHC)

Xenograft tumors were harvested 6 hours after last treatment with rapamycin or INK128, and 24 hours after last treatment with ganetespib. Resected tumors were used for immunoblotting and/or used for immunohistochemistry (IHC). For IHC tumors were fixed with 10% formalin overnight and stored in 70% ethanol solution until paraffin embedding. Paraffin-embedded sections, both unstained and stained (hematoxylin and eosin stain), were prepared by the Rodent Histopathology Core, Harvard Medical School.

For IHC, slides were rinsed with a series of xylene, ethanol, 95% ethanol, 80% ethanol, and PBS. To unmask the epitopes slides were boiled in 10 mM sodium citrate for 60 minutes and cooled to room temperature. Slides were washed with PBS. Slides were put into peroxidase blocking reagent (225 mL methanol, 25 mL 30% hydrogen peroxide) for 10 minutes, and washed with PBS. Afterward, slides were washed with distilled water, sections were counterstained with hematoxylin and washed under running water.

Cell proliferation was assessed using an antibody against PCNA and the HistoMouse kit-plus kit (Invitrogen/Thermo Fisher Scientific), using AEC single solution chromogen, and then counterstained with hematoxylin. Sections were mounted with cover glass and Fluoromount G media.

For TSC2 and pS6 IHC primary antibodies against pS6 (Ser235/236) (2211, Cell Signaling Technology) at a dilution of 1:250 and TSC2 (sc-271314, Santa Cruz Biotechnology) at a dilution of 1:100 were used and incubated at 4 °C overnight, rinsed, and then treated with anti-rabbit-HRP secondary antibody at room temperature for 60 minutes, followed by rinsing and slide preparation as above.

To identify apoptotic cells, ApopTag plus peroxidase in situ apoptosis detection kit (Merck Millipore) was used according to the manufacturer’s instructions.

### Statistical analysis

The quantitative data of the xenograft tumor experiments are reported as the mean ± standard deviation for at least 5 tumors. Tumor sizes for the different treatment groups were compared using the Wilcoxon Rank Sum test using GraphPad Prism software. P-values less than 0.05 were considered statistically significant.

## Results

TSC1/TSC2 are known recessive oncogenes whose loss is known to activate mTORC1. Here we sought to identify and characterize cancer cell lines with complete loss of either TSC1 or TSC2, and then to examine their sensitivity to a broad panel of kinase inhibitors, with a goal to identify additional inhibitors beyond rapalogs.

### Identification of cancer cell lines lacking expression of either TSC1 or TSC2

Data from the Broad Institute Cancer Cell Line Encyclopedia (CCLE) (26) was reviewed to identify tumor cell lines with mutations in either *TSC1* or *TSC2*. Among 1457 cell lines, 49 were reported to have mutations in *TSC1* and 111 to have mutations in *TSC2*. These mutations were reviewed to identify those with nonsense mutations, frameshift deletions or insertions, or in-frame deletions in either *TSC1* or *TSC2*, yielding 4 cell lines with probable mutations in *TSC1* and 10 with probable mutations in *TSC2* (**Table 1**).

**Table 1.** CCLE cell lines with reported nonsense mutations, frameshift deletions or insertions, or in-frame deletions in TSC1 and TSC2 (adapted from 27, 28)

| tumor sample | allele ratio | genome change, hg19 | variant classification | variant type | reference allele | tumor seq. allele | cDNA change | codon change | protein change |
| --- | --- | --- | --- | --- | --- | --- | --- | --- | --- |
| ***TSC1*, chromosome 9** |  |  |  |  |  |  |  |  |  |
| PEER_HAEM_AND_LYMPH | 1.00 | g.chr9:135798751C>T | Nonsense_Mutation | SNP | C | T | c.492G>A | c.(490-492)TGG>TGA | p.(Trp164*) |
| KM12_LARGE_INTESTINE | 0.56 | g.chr9:135797261_135797261delA | Frame_Shift_Del | DEL | A | - | c.608_608delT | c.(607-609)TTGfs | p.(Leu203Cysfs*7) |
| 2313287_STOMACH | 0.52 | g.chr9:135785954G>A | Intron | SNP | G | A | c.1263+4C>T |  |  |
| CW2_LARGE_INTESTINE | 0.47 | g.chr9:135781157_135781157delG | Frame_Shift_Del | DEL | G | - | c.1808_1808delC | c.(1807-1809)CCGfs | p.(Pro603Argfs*26) |
| ***TSC2*, chromosome 16** |  |  |  |  |  |  |  |  |  |
| NCIH1651_LUNG | 0.33 | g.chr16:2100401G>T | Nonsense_Mutation | SNP | G | T | c.139G>T | c.(139-141)GAA>TAA | p.(Glu47*) |
| CW2_LARGE_INTESTINE | 0.60 | g.chr16:2108787_2108788insT | Frame_Shift_Ins | INS | - | T | c.888_889insT | c.(886-891)GTGTTTfs | p.(Val299Cysfs*39) |
| HEC151_ENDOMETRIUM | 0.42 | g.chr16:2121790_2121791insAG | Frame_Shift_Ins | INS | - | AG | c.1952_1953insAG | c.(1951-1953)CCAFs | p.(Gly654Glufs*45) |
| DV90_LUNG | 0.36 | g.chr16:2121791_2121792delAG | Frame_Shift_Del | DEL | AG | - | c.1953_1954delAG | c.(1951-1956)CCAGAGfs | p.(Gly654Leufs*2) |
| OVK18_OVARY | 0.67 | g.chr16:2121791_2121792delAG | Frame_Shift_Del | DEL | AG | - | c.1953_1954delAG | c.(1951-1956)CCAGAGfs | .(Gly654Leufs*2) |
| MFE319_ENDOMETRIUM | 0.42 | g.chr16:2122880C>T | Nonsense_Mutation | SNP | C | T | c.2251C>T | c.(2251-2253)CGA>TGA | p.(Arg751*) |
| SNU-886_LIVER | 0.97 | g.chr16:2129160C>T | Nonsense_Mutation | SNP | C | T | c.3094C>T | c.(3094-3096)CGA>TGA | p.(Arg1032*) |
| SNU-878_LIVER | 1.00 | g.chr16:2134999C>G | Nonsense_Mutation | SNP | C | G | c.4541C>G | c.(4540-4542)TCA>TGA | p.(Ser1514*) |
| HEC151_ENDOMETRIUM | 0.45 | g.chr16:2138565G>A | Missense_Mutation | SNP | G | A | c.5378G>A | c.(5377-5379)CGG>CAG | p.(Arg1793Gln) |
| HCC95_LUNG | 0.60 | g.chr16:2138578_2138580delCTC | In_Frame_Del | DEL | CTC | - | c.5391_5393delCTC | c.(5389-5394)ATCTCC>ATC | p.(Ser1799del) |

All of these cell lines were obtained, and mutations were confirmed by Sanger sequencing (**S1 Fig**). The cell line PEER showed a homozygous nonsense mutation in *TSC1*; SNU-878 and SNU-886 showed homozygous nonsense mutations in *TSC2*. All other cell lines had mutations with allele ratios around 0.5.

### mTOR pathway assessment

As further confirmation of mutational status for these cell lines, we performed immunoblotting to assess expression of TSC1 and TSC2, and mTORC1 signaling, which should be activated in the absence of either TSC1 or TSC2 (**Fig 1**). There was no expression of TSC1 in PEER cells, and absence of TSC2 in SNU-878 and SNU-886 cells. All other cell lines showed some degree of expression of TSC1 or TSC2, as expected. Levels of AKT, S6K, and S6 were similar among all cell lines. Persistent activation of S6K by phosphorylation at Thr389, and S6 by phosphorylation at Ser240/244 in the absence of serum (29, 30), was seen in PEER, SNU-878, and SNU-886, consistent with constitutive mTORC1 activation. Increased pS6K (Thr389) and pS6 (Ser240/244) levels in the absence of serum were also seen in MFE-319 and OVK18 cells, both of which are known to have PTEN mutations (26, 31, 32).

**Fig 1.**
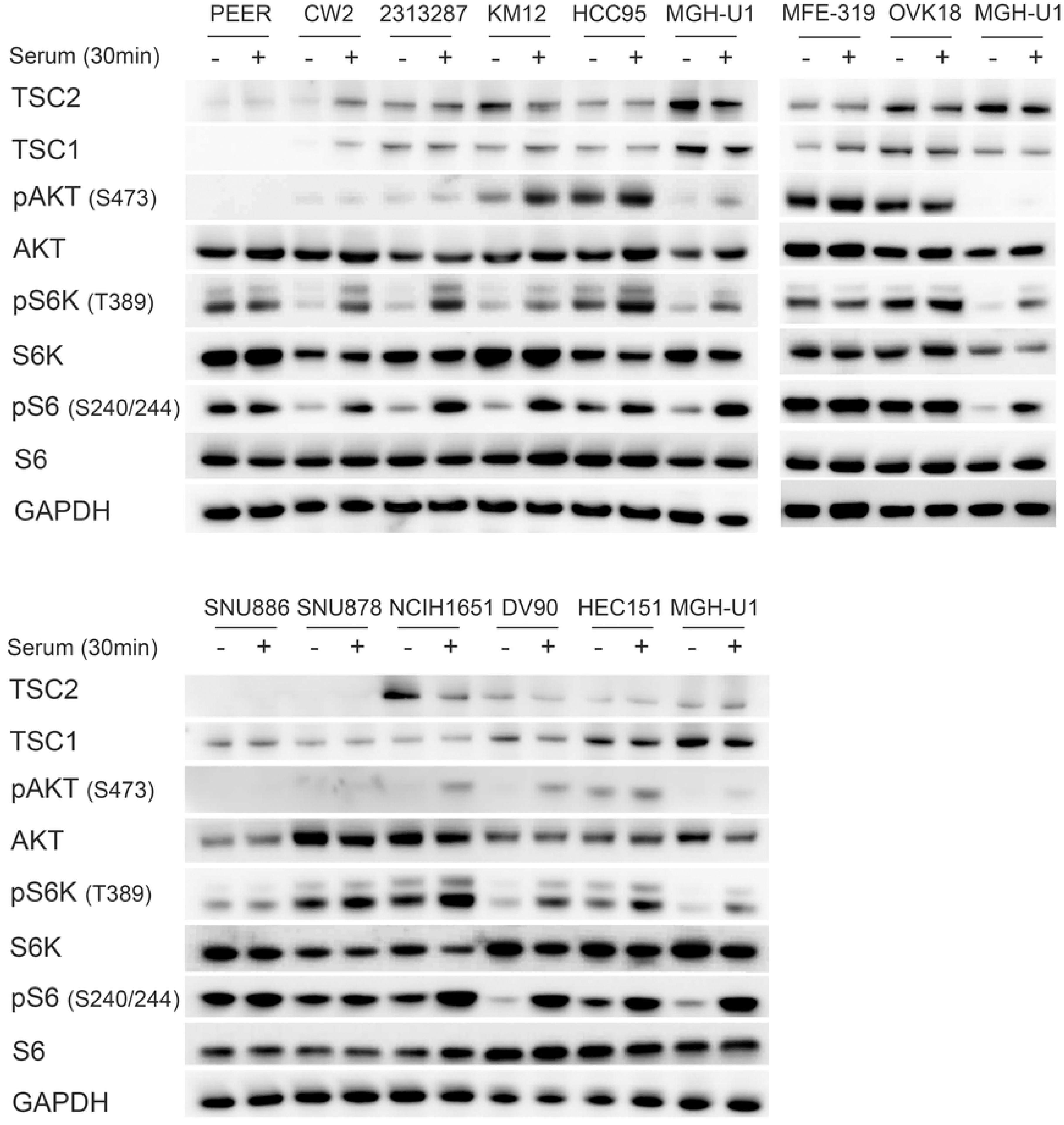
Identification and characterization of cell lines with *TSC1* or *TSC2* mutations. Cells were serum starved for 24h (−) or had serum add back for 30 min (+) after starvation. MGH-U1, a bladder cancer cell line with normal mTOR signaling, was used as a control.

Absence of TSC1 expression was seen in the cell line PEER and absence of TSC2 expression in the cell lines SNU-878 and SNU-886. Constitutive mTOR activity was seen, indicated by increased pS6K (Thr389) and pS6 (Ser240/244) expression in serum starvation, in PEER, MFE-319, OVK18, SNU-886, and SNU-887 cells. GAPDH was used as a loading control.

425 cancer cell lines were also screened for large deletions in the *TSC1* and *TSC2* genes by the MLPA method (see Methods). After a round of screening followed by repeat assessment using probes for each exon of *TSC1* and *TSC2*, the osteosarcoma cell line CAL-72 showed homozygous deletion of exons 6 to 23 of *TSC1*; and the hepatocellular carcinoma cell line SNU-398 showed homozygous deletion of exons 10-41 of *TSC2*.

Hence the five cell lines, PEER, CAL-72, SNU-878, SNU-886, and SNU-398, were studied in greater detail. CAL-72 and PEER cells both showed complete loss of TSC1 and reduced TSC2 expression (**Fig 2**), consistent with the effect of TSC1 in stabilizing TSC2 protein levels through heterocomplex formation, as shown previously (33). SNU-878, SNU-886, and SNU-398 showed complete loss of TSC2 and normal TSC1 expression (**Fig 2**). The levels of mTOR, AKT, BIP, S6K, and S6 were similar among all cell lines. pS6K (Thr389), pS6 (Ser235/236 and Ser240/244), and pEIF2α (Ser51) were increased in the absence of serum in all TSC1 or TSC2 deficient cell lines, indicating constitutive mTORC1 activation. The control cell line MGH-U2 showed low levels of pS6K (Thr389) and pS6 (Ser235/236 and Ser240/244) when serum-starved, indicating normal mTORC1 regulation.

**Fig 2.**
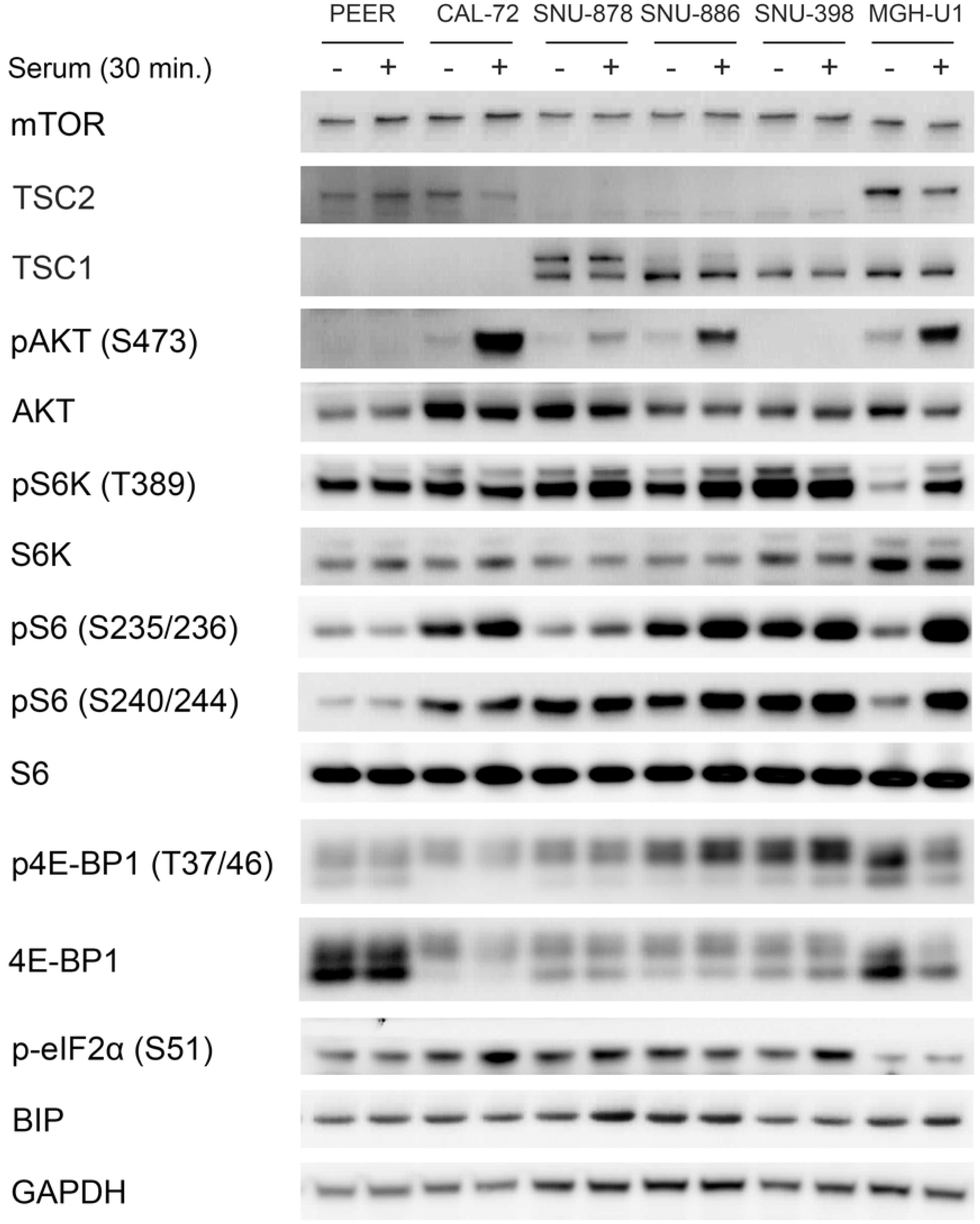
Characterization of TSC1 and TSC2 deficient cell lines. Immunoblots of TSC1 and TSC2 deficient cell lines were performed to examine multiple components of the mTORC1 signaling pathway. Cells were serum starved for 24h (−) or had serum add back for 30 min (+) after starvation. CAL-72 and PEER cells showed no TSC1 and reduced TSC2 expression. SNU-878, SNU-886, and SNU-398 showed an absence of TSC2 expression. pS6K (Thr389), pS6 (Ser235/236 and Ser240/244) p4E-BP1 (Thr37/46), and pEIF2 (Ser51) showed a strong expression in a serum starved condition in all TSC1 or TSC2 deficient cell lines, indicating constitutive mTORC1 activation. GAPDH and BIP were used as loading controls.

PEER and SNU-398 showed no AKT activation, as assessed by lack of pAKT (Ser473), in serum absence and serum add back conditions (**Fig 2**). This has been observed previously in many other cell lines with a constitutive mTORC1 activation, consistent with an active feedback suppression due to mTORC1 activation (30, 34, 35). CAL-72, SNU-886, SNU-878, and the control cell line showed increased pAKT-Ser473 levels in response to serum add back.

Variable 4E-BP1 expression was seen among these cell lines, but all TSC1 or TSC2 deficient cell lines showed similar levels of expression of p4E-BP1-Thr37/Thr46 in serum-deprived and serum add back conditions. Expression of p-eIF2α (Ser51) was also higher in TSC1 or TSC2 deficient cells in comparison to the control cell line (**Fig 2**).

### Kinase inhibitor library screen and IC_50_ determination

The cell lines SNU-878, SNU-886, SNU-398, CAL-72, and PEER were screened for sensitivity to kinase inhibition using a kinase inhibitor-focused library (LINCS), which contained 197 selective kinase inhibitors (**S1 Table**). This library was composed of commercially available and self-developed pharmacophore-diverse ATP competitive kinase inhibitors. The library contained inhibitors against multiple different kinases, including those involved in: regulation of the cell cycle (cyclin-dependent kinases (CDKs), CLK2, Polo-like kinases (PLKs), and Aurora kinases); DNA damage repair (checkpoint kinases (CHKs), CNSK1E, and ATMs); ligand - receptor tyrosine kinase signaling (VEGFR, HER2, EGFR, PDGFR, and FGFR); mitogen-activated protein kinases (MAPKs, MEKs, ERKs, B-Raf, mTOR, PI3Ks, AKT); intracellular tyrosine kinases (FLT3, Srcs, Syk, FAKs, BTK/BMX, c-Kit or c-Met, c-Raf, ABL, Bcr-Abl, JAKs, LCK, Tie2, DDR, RET); serine/threonine kinases (Rock 2, Rsk2, DNA-PKcs, BRSK2, RIPK1, PKC, TBK1, MNK2, Wee1 and LOKs); and a further diverse set (GSK3, PDK1, Alk, IKK-2, ITK, IRAK1, CSF1R, ULK1, LRRK2, EPHB3, CAMKs, PIKfyve, IGF-1R, PARPs, HSP-90, p53, Rho, Bcl-2, c-FMS, PI4KIII, EPHB4 and CK1). In this initial screen, a single dose of inhibitor was used, 600nM. All compounds that showed > 50% reduction in growth for one or more cell lines in this assay were considered initial positive hits and were subject to further study on all cell lines. A wide variety of inhibitors showed a positive signal in this initial assay, including multiple mTOR, CDK, PLK, CHK, Aurora, and HSP90 inhibitors (**Table 2**).

**Table 2.**
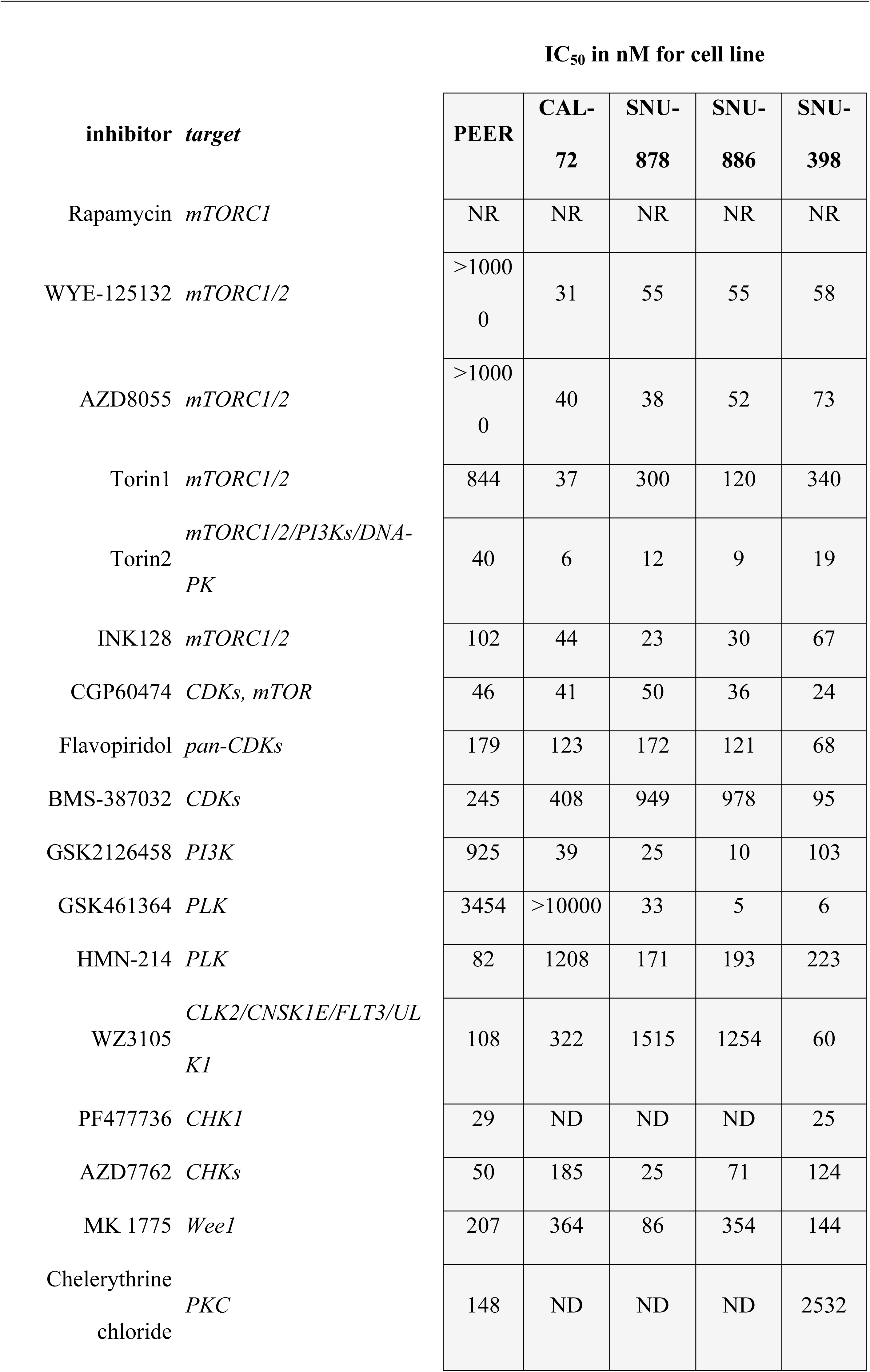
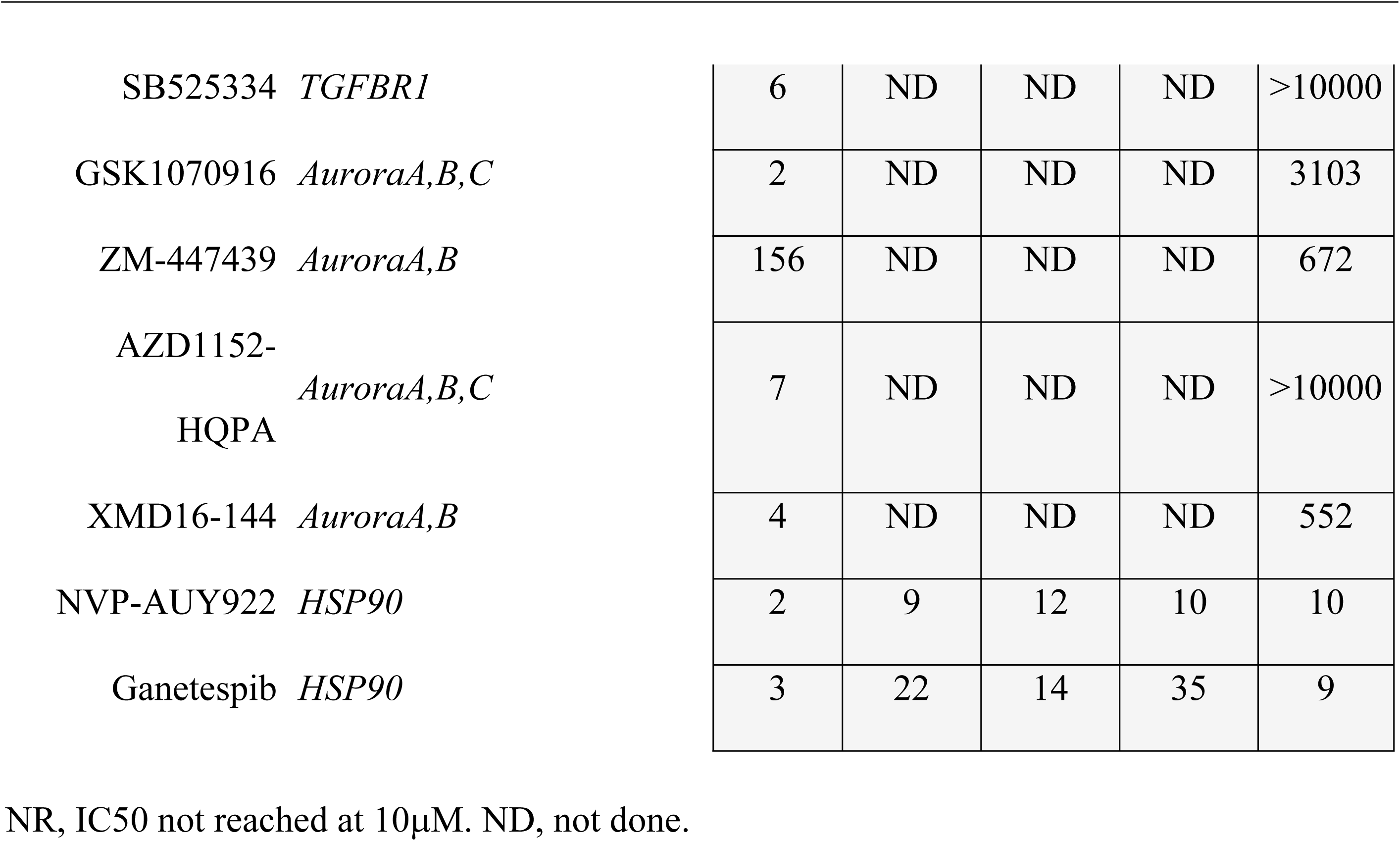
IC_50_ determination of the most effective drugs.

### mTOR and PI3K inhibitor effects on TSC1/TSC2 null cell lines

Candidate inhibitors were studied against most or all of the TSC1/TSC2-deficient cell lines in standard 96 well plate growth assays over a dose range of 1 nM to 10μM (**Table 2**). We focus on the results with the mTOR inhibitors first. Rapamycin, an mTORC1 allosteric inhibitor, did not achieve IC50 over this dose range (**S2 Fig**). Among the 5 other mTOR inhibitors, Torin2 showed the lowest IC50 for each of the 5 cell lines (**S3-S7 Fig**), similar to previous studies (39). This may relate to its intrinsic potency, or to its additional inhibitory effects on DNA-PKcs, PIK3C, PIK3R, and PI4KB (36). The other mTOR inhibitors studied also have inhibitory effects on other kinases (**S1 Table**) (37-39). Among the cell lines, PEER was universally the most resistant, requiring the highest dose of all inhibitors to achieve IC50 (**Table 2, S3-S7 Fig**).

### Cell cycle inhibitor effects on the TSC1/TSC2 null cell lines

Several cell cycle inhibitors showed significant growth inhibition for the TSC mutant cells lines, including CDK, PLK, and Aurora kinase inhibitors (40). However, these inhibitors in general also have cross-reactivity with several other kinases. All cell lines were sensitive to Flavopiridol with fairly uniform IC_50_ values ranging from 68 to 179 nM (**Table 2**), and were even more sensitive to CGP60474, which may have been due to inhibitory effects on mTOR (**Table 2, S8-10 Fig**).

The three *TSC2* null liver carcinoma cell lines SNU-886, SNU-878, and SNU-398 were uniformly sensitive to GSK461364, a PLK1 inhibitor, while in contrast the *TSC1* null cell lines PEER and CAL-72 were not sensitive at all (**Table 2**) with IC_50_ values ranging from 5 to 33 nM. PLK1 promotes G2/M-phase transition (41) and PLK1 inhibition stimulates lysosomal localization of mTOR and consequently decreased autophagy activation (42). The SNU cell lines were less sensitive to another PLK inhibitor, HMN-214, an oral pro-drug of HMN-176, which interferes with the location and consequently the function of PLK1 (43) (**Table 2, S11** and **S12 Fig**).

*TSC1* null PEER cells were very sensitive to all Aurora inhibitors with IC50 values ranging from 2 to 156 nM (**Table 2**). In contrast, *TSC2* null SNU-398 cells were much less sensitive to Aurora inhibitors (**Table 2, S13-S16 Fig**).

### HSP90 inhibitors

All cell lines were very sensitive to both NVP-AUY922 and ganetespib, HSP90 inhibitors, with IC50 values ranging from 2 to 12 nM and 3 to 35 nM, respectively (**Table 2, S17** and **S18 Fig**).

### Combination treatment with mTORC1 and HSP90 inhibitors

Given the activity of mTORC1 inhibitors and HSP90 inhibitors individually, we examined the potential synergistic effect of combination treatment using drugs from both classes. Ganetespib was serially diluted three-fold from 10 μM to 1.5 nM. The Torin2 concentration was fixed at 5nM, and the rapamycin concentration at 20 nM. Combined therapy of genetespib and rapamycin or genetespib and Torin2 showed some degree of additive effect with lower IC_50_ values in some cases (**Table 3, Fig 3**). However, the significance of this additive effect was not clear.

**Table 3.** IC_50_ determination of Torin2, Ganetespib and combination therapy.

|  |  | IC<sub>50</sub> in nM for cell line |  |  |  |  |
| --- | --- | --- | --- | --- | --- | --- |
| inhibitor | target | PEER | CAL-72 | SNU-878 | SNU-886 | SNU-398 |
| Torin2 | *mTORC1/2/PI3Ks/DNA-PK* | 40 | 6 | 12 | 9 | 19 |
| Ganetespib | *HSP90* | 3 | 22 | 14 | 35 | 9 |
| Ganetespib + Torin2 (fixed at 5nM) |  | 6 | 20 | 8 | 9 | 4 |
| Ganetespib + Rapamycin (fixed at 20nM) |  | 6 | 17 | 6 | 5 | 2 |

**Fig 3.**
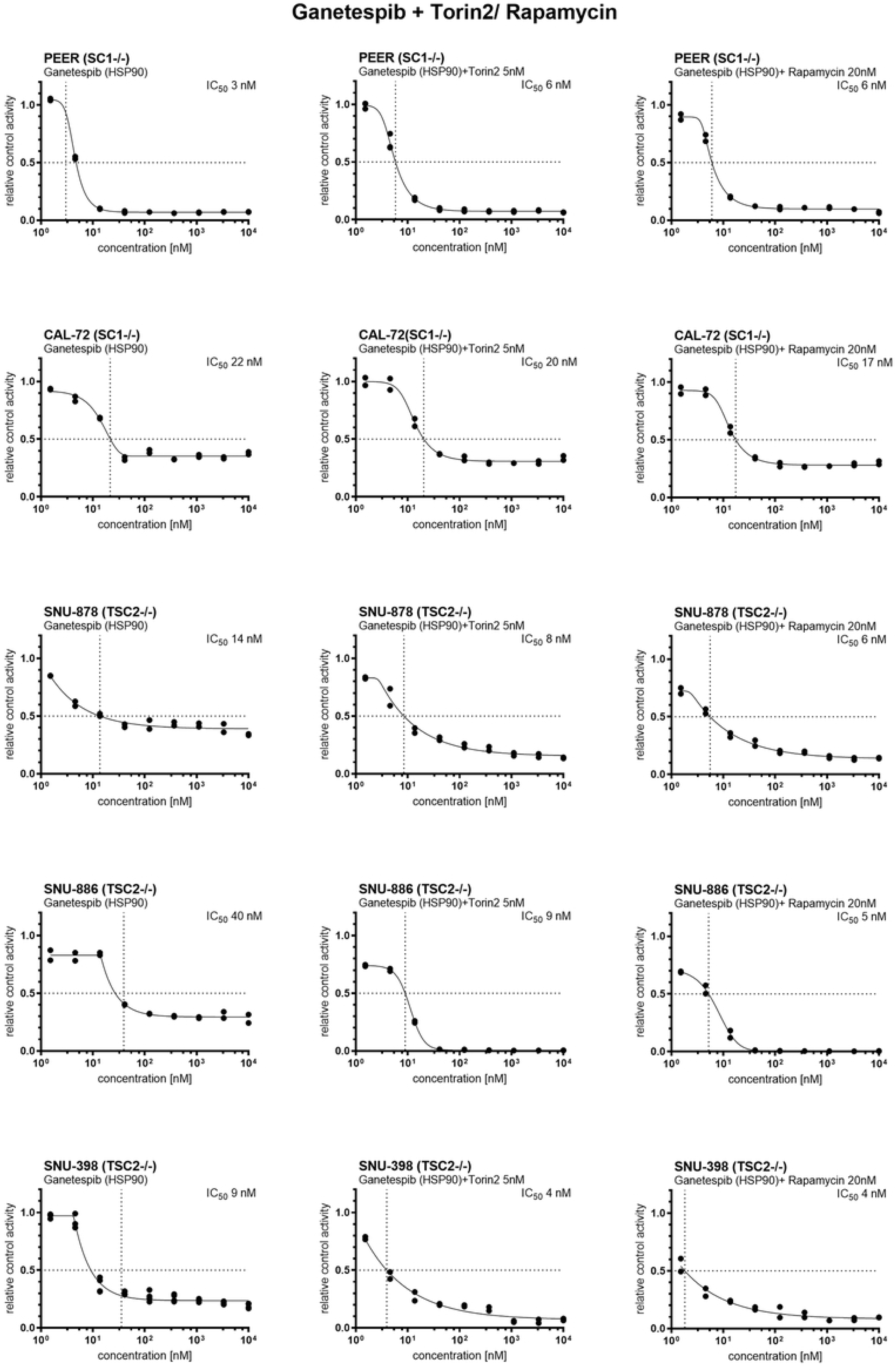
Cell viability after ganetespib, ganetespib with Torin2 and ganetespib with rapamycin treatment. Ganetespib was serially diluted three-fold from 10 μM to 1.5 nM. The Torin2 concentration was 5nM, and the rapamycin concentration 20 nM. Cell viability was determined 72h after treatment using CellTiter-Glo and XLfit4.0 software. Cell viability is shown in relative control activity, n= 2-4. IC_50_ was calculated as the drug concentration that reduced cell viability by 50% compared to untreated cells. All TSC1 or TSC2 deficient tumor cell lines showed a strongly decreased cell viability after treatment with ganetespib. Combined therapy of genetespib and rapamycin or genetespib and Torin2 showed an additive effect with even lower IC50 values. However, the significance of this additive effect was not clear.

### Generation and characterization of add back derivatives of the TSC2 mutant cell lines

TSC2-encoding or empty vectors were added via retroviral transfection to the TSC2-/- cell lines SNU-886, SNU-878, and SNU-398, to generate TSC2 add-back and control cell lines. Immunoblot analysis demonstrated that TSC2 expression was restored in the TSC2 add-back derivatives. pS6 (Ser240/244) levels were decreased in the absence of serum conditions in the TSC2 add back cells, indicating return to normal regulation of mTORC1. pAKT (Ser473) expression was quite low in the TSC2 add back cells, but increased significantly in response to serum stimulation in the cell lines SNU-886 and SNU-878, compared to the levels of native or empty vector-transfected cells. SNU-398 did not show a pAKT (Ser473) expression increase in TSC2 add back cells (**Fig 4**).

**Fig 4.**
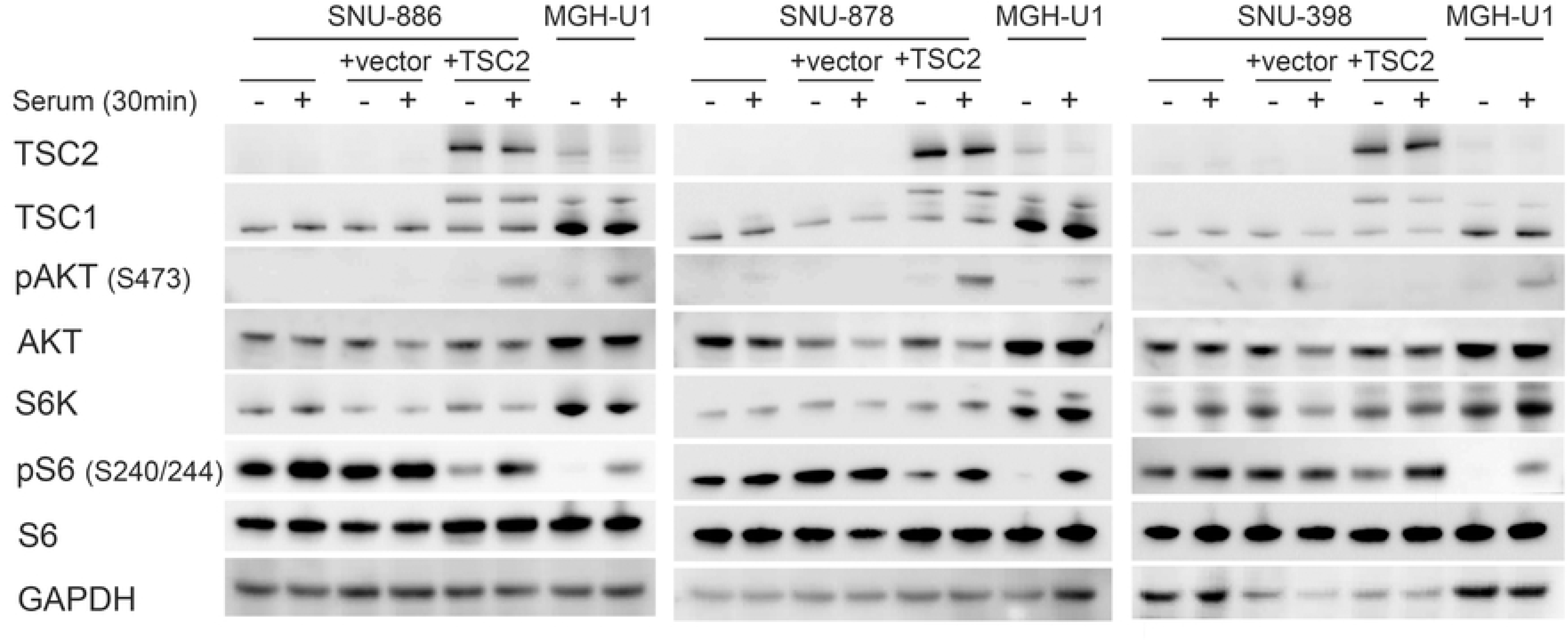
Expression and signaling effects of TSC2 add back to TSC2 deficient cells. *TSC2* cDNA or an empty vector was added to TSC2 deficient cell lines via retroviral transfection. Cells were serum starved for 24h (−) or had serum add back for 30 min (+). In the TSC2 add back cells there is a strong TSC2 expression, a decreased pS6 (S240/244) expression in the absence of serum, and increased pAKT (Ser473) expression following serum add back. No pAKT (Ser473) was seen in SNU-398 cells, with or without TSC2 addback.

These three TSC2 add back cell lines were used in drug testing experiments similar to what was done above. Interestingly, TSC2 add back cells had a 3 to 6-fold higher IC50 for both Torin1 and INK 128 than their empty vector controls or original unmanipulated cells (**Table 4, S5** and **S7 Fig**). However, in contrast, the three other mTOR kinase inhibitors, WYE-125132, Torin2, and AZD8055, showed no significant difference in IC50 between TSC2 add back cells and cells with EV or the unmanipulated starting cells (**Table 4, S3, S4, S6 Fig**). Minor differences in IC50 was seen for these cell lines using multiple other inhibitors (**Table 4, S2, S8-S12, S15, S17-S21 Fig)**.

**Table 4.**
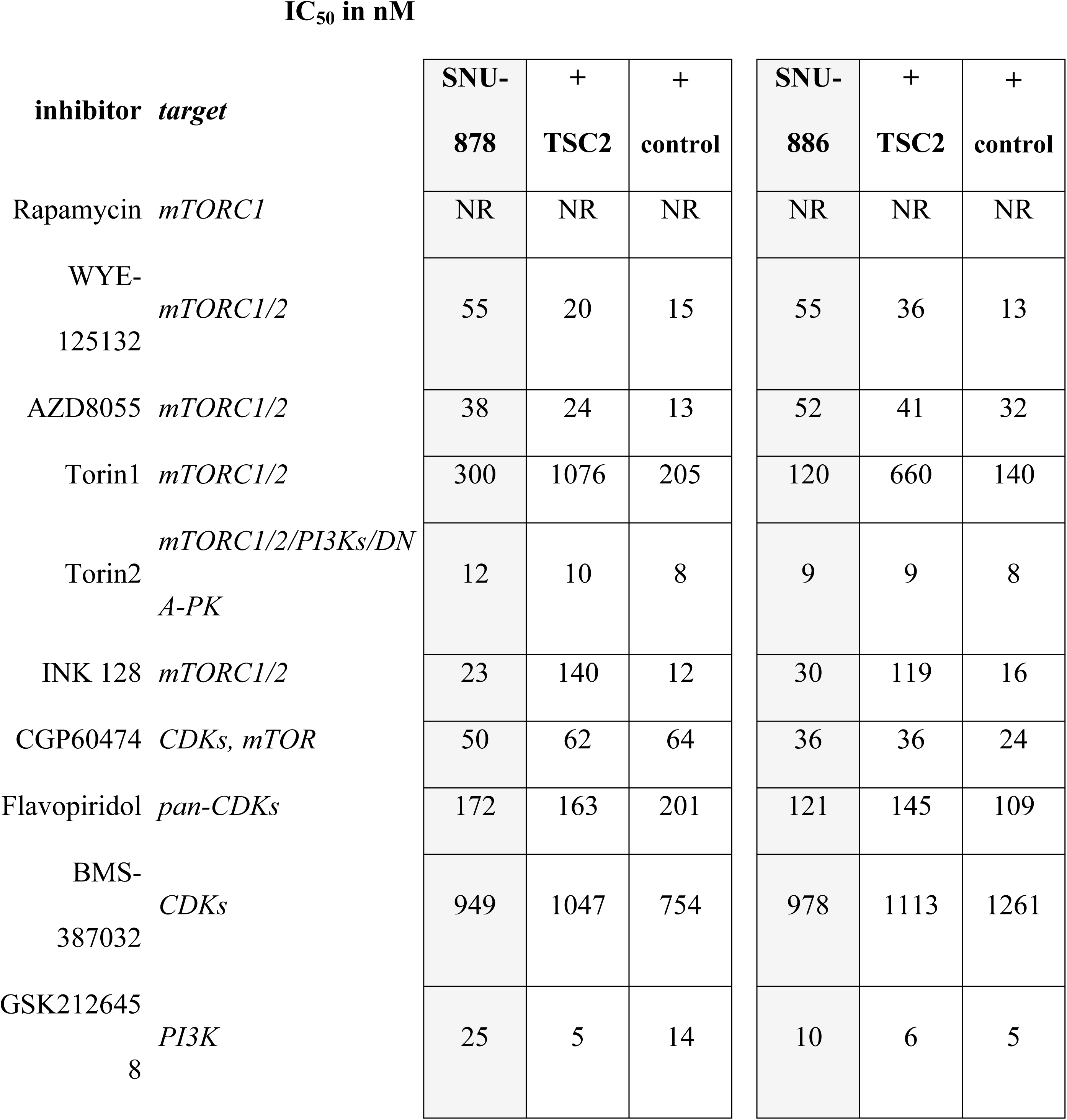
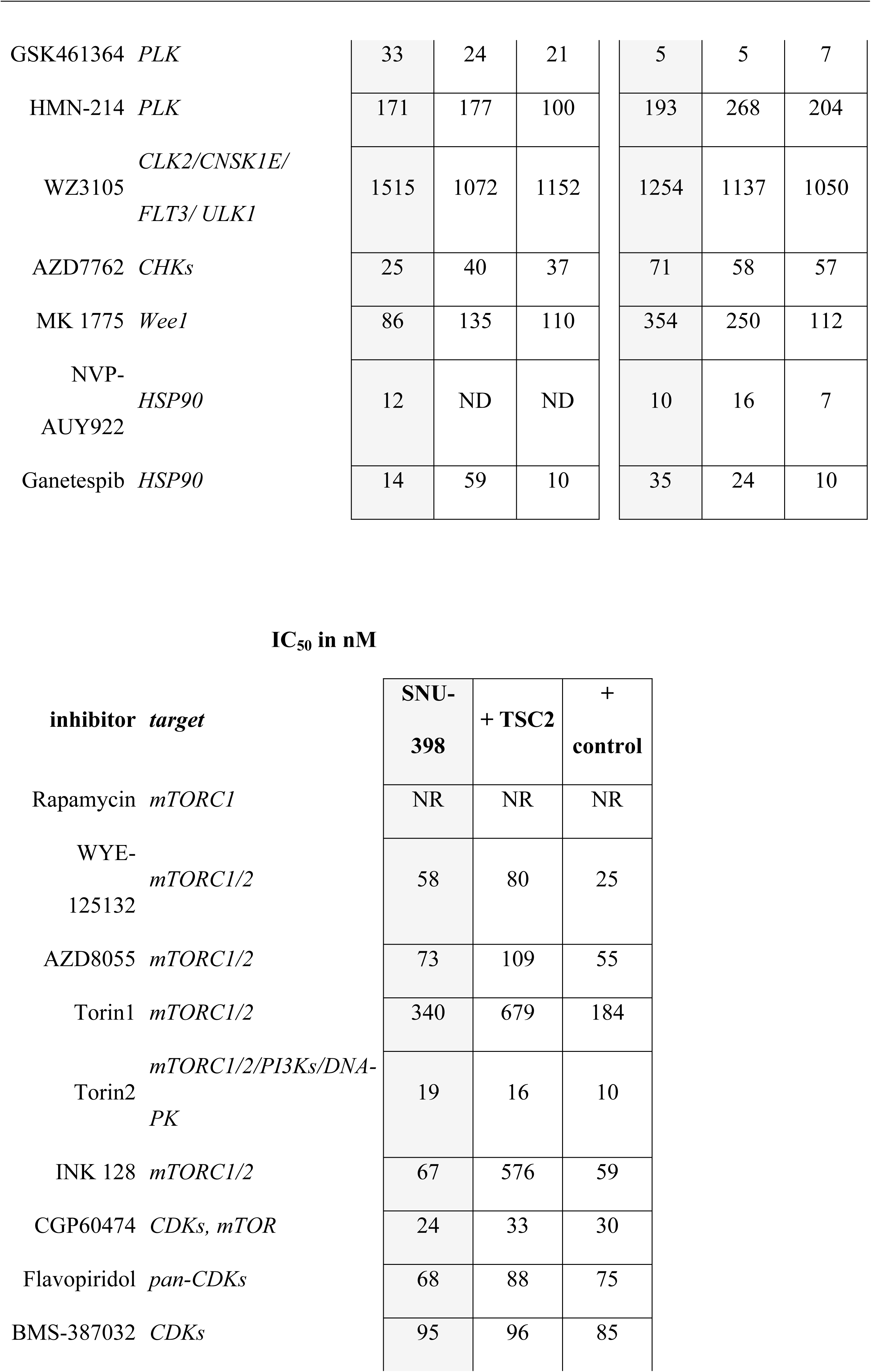
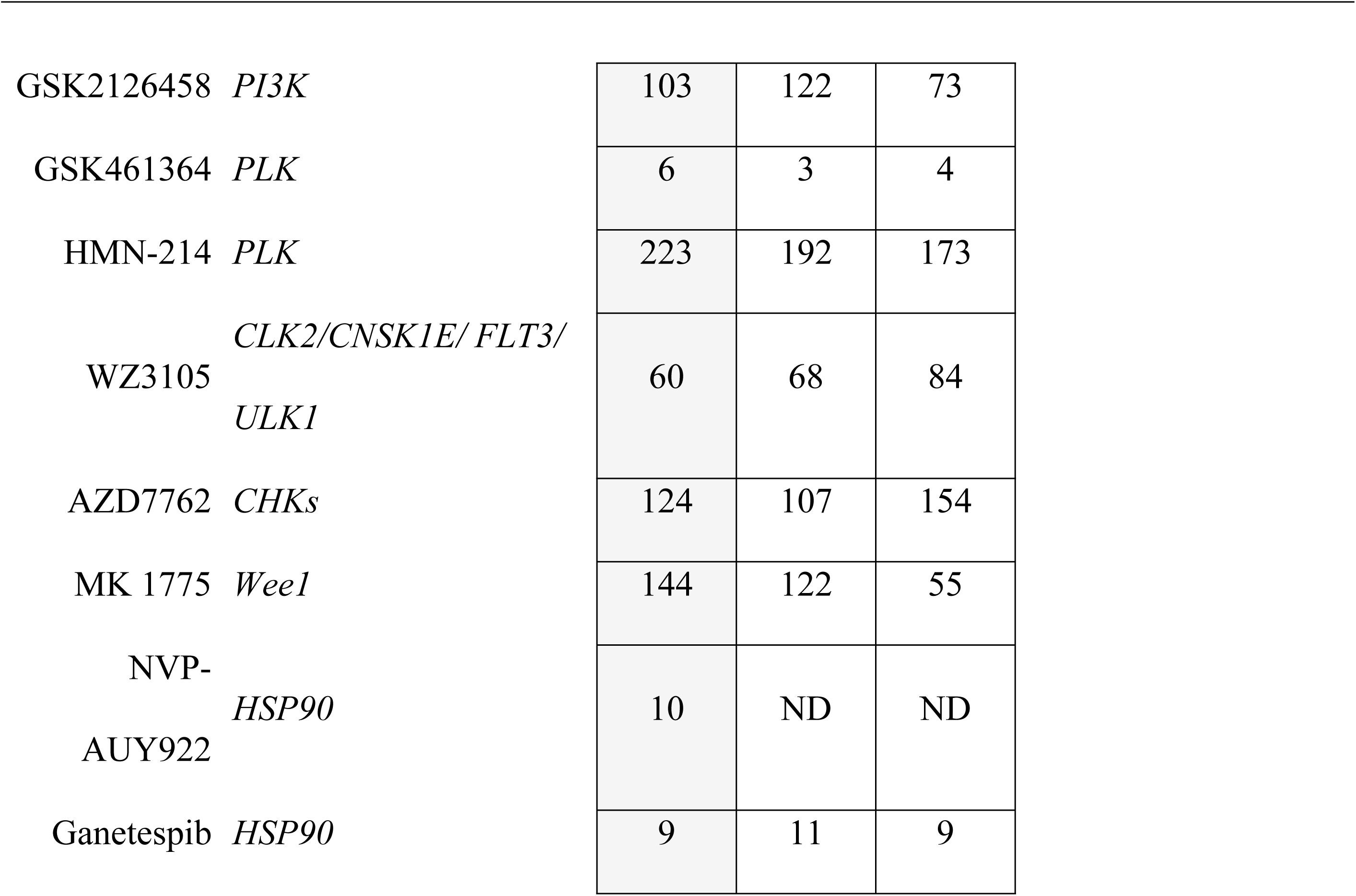
IC_50_ determination of TSC2 deficient cells, cells with TSC2 add back and control vector.

|  |  | IC<sub>50</sub> in nM |  |  |  |  |  |
| --- | --- | --- | --- | --- | --- | --- | --- |
| inhibitor | target | SNU-878 | + TSC2 | + control | SNU-886 | + TSC2 | + control |
| Rapamycin | *mTORC1* | NR | NR | NR | NR | NR | NR |
| WYE-125132 | *mTORC1/2* | 55 | 20 | 15 | 55 | 36 | 13 |
| AZD8055 | *mTORC1/2* | 38 | 24 | 13 | 52 | 41 | 32 |
| Torin1 | *mTORC1/2* | 300 | 1076 | 205 | 120 | 660 | 140 |
| Torin2 | *mTORC1/2/PI3Ks/DNA-PK* | 12 | 10 | 8 | 9 | 9 | 8 |
| INK 128 | *mTORC1/2* | 23 | 140 | 12 | 30 | 119 | 16 |
| CGP60474 | *CDKs, mTOR* | 50 | 62 | 64 | 36 | 36 | 24 |
| Flavopiridol | *pan-CDKs* | 172 | 163 | 201 | 121 | 145 | 109 |
| BMS-387032 | *CDKs* | 949 | 1047 | 754 | 978 | 1113 | 1261 |
| GSK2126458 | *PI3K* | 25 | 5 | 14 | 10 | 6 | 5 |

|  |  |  |  |  |  |  |
| --- | --- | --- | --- | --- | --- | --- |
| GSK461364 *PLK* | 33 | 24 | 21 | 5 | 5 | 7 |
| HMN-214 *PLK* | 171 | 177 | 100 | 193 | 268 | 204 |
| WZ3105 *CLK2/CNSK1E/*<br>*FLT3/ULK1* | 1515 | 1072 | 1152 | 1254 | 1137 | 1050 |
| AZD7762 *CHKs* | 25 | 40 | 37 | 71 | 58 | 57 |
| MK 1775 *Wee1* | 86 | 135 | 110 | 354 | 250 | 112 |
| NVP-AUY922 *HSP90* | 12 | ND | ND | 10 | 16 | 7 |
| Ganetespib *HSP90* | 14 | 59 | 10 | 35 | 24 | 10 |

IC<sub>50</sub> in nM
| **inhibitor target** | **SNU-398** | **+ TSC2** | **+ control** |
| --- | --- | --- | --- |
| Rapamycin *mTORC1* | NR | NR | NR |
| WYE-125132 *mTORC1/2* | 58 | 80 | 25 |
| AZD8055 *mTORC1/2* | 73 | 109 | 55 |
| Torin1 *mTORC1/2* | 340 | 679 | 184 |
| Torin2 *mTORC1/2/PI3Ks/DNA-PK* | 19 | 16 | 10 |
| INK 128 *mTORC1/2* | 67 | 576 | 59 |
| CGP60474 *CDKs, mTOR* | 24 | 33 | 30 |
| Flavopiridol *pan-CDKs* | 68 | 88 | 75 |
| BMS-387032 *CDKs* | 95 | 96 | 85 |

|  |  |  |  |  |
| --- | --- | --- | --- | --- |
| GSK2126458 | *PI3K* | 103 | 122 | 73 |
| GSK461364 | *PLK* | 6 | 3 | 4 |
| HMN-214 | *PLK* | 223 | 192 | 173 |
| WZ3105 | *CLK2/CNSK1E/FLT3/ULK1* | 60 | 68 | 84 |
| AZD7762 | *CHKs* | 124 | 107 | 154 |
| MK 1775 | *Wee1* | 144 | 122 | 55 |
| NVP-AUY922 | *HSP90* | 10 | ND | ND |
| Ganetespib | *HSP90* | 9 | 11 | 9 |

### Effects of rapamycin, Torin1 and ganetespib on protein expression and mTOR signaling

To examine the impact of these drugs on mTOR signaling in the TSC-mutant cell lines, PEER, CAL-72, SNU-886, SNU-878, and SNU-398 cells were treated with rapamycin 10 nM, Torin1 250 nM, and ganetespib at doses ranging from 100 nM to 1 μM for 24 hours. Both rapamycin and Torin1 had little effect on expression levels of mTOR, AKT, S6K, and S6; while both reduced expression of pS6K (T389) and pS6 (S240/244) to a major degree in the absence of serum, as well as with serum stimulation (**Fig 5**). These findings were expected given their inhibition of the constitutively activated mTORC1. pAKT (Ser473) levels were also lower in Torin1-treated cells compared to rapamycin-treated cells, as expected given its inhibition of mTORC2 as well as mTORC1. pAKT (Ser473) levels were higher in rapamycin-treated SNU-886 and SNU-878 cells than controls, due to feedback activation of IGF1 and hence mTORC2, which phosphorylates AKT, as described previously (35, 49) PEER and SNU-398 did not show pAKT (Ser473) expression regardless of treatment, suggesting that other mutations and/or pathways effects were operative in these cells (**Fig 5**).

**Fig 5.**
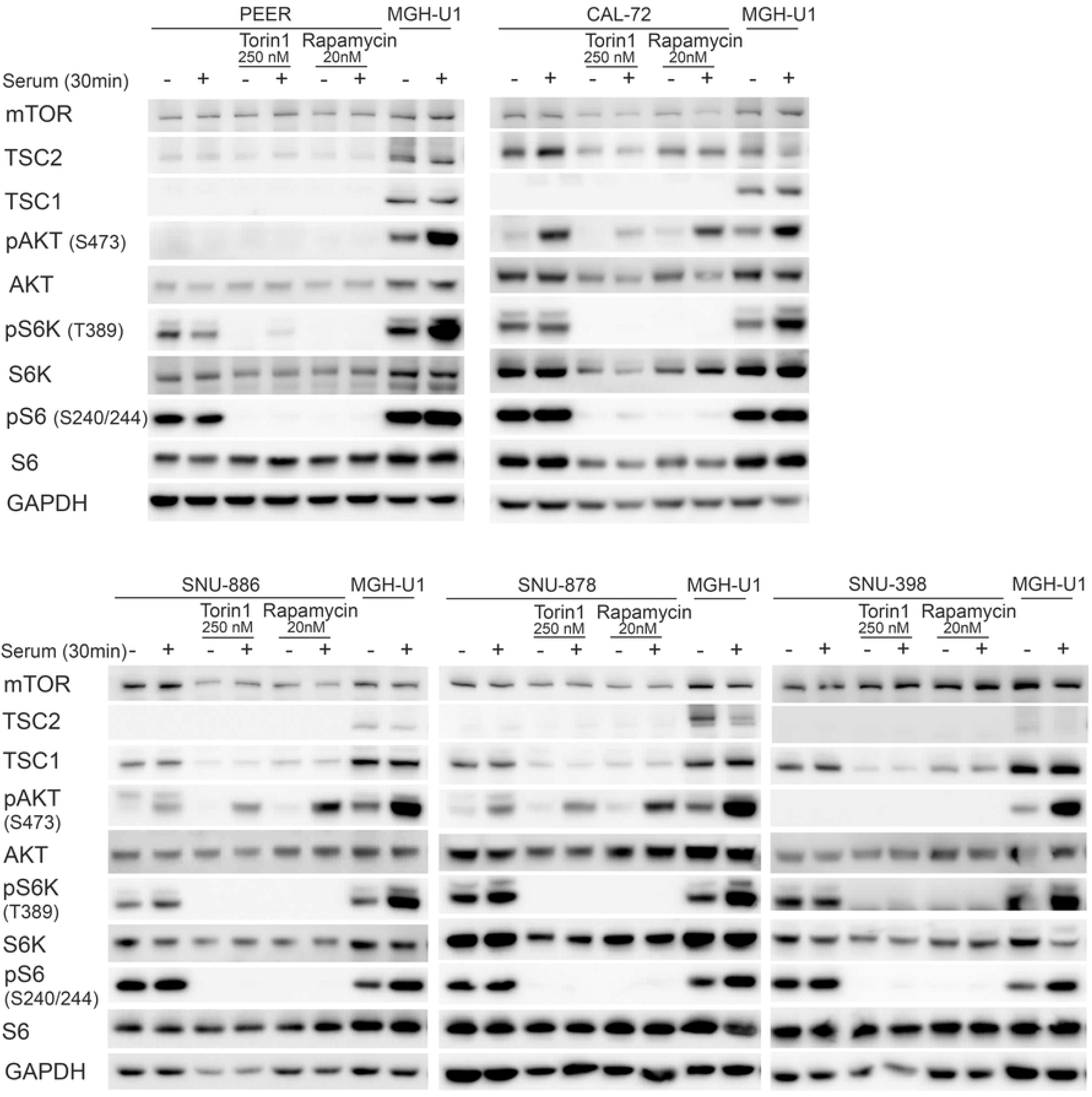
Impact of rapamycin and Torin1 on constitutively activated mTOR signaling pathway in TSC1 or TSC2 deficient cells. Cells were treated with Torin1 (250 nM) or rapamycin (20 nM) for 24h. Untreated cells and MGH-U1 were used as controls. Cells were serum starved for 24h (−) or received after starvation serum add back for 30 min (+). Rapamycin and Torin1-treated cells show no expression of pS6K (Thr389) and pS6 (Ser 240/244) in serum absence or stimulated conditions compared to the upregulated expression in untreated cells. pAKT (Ser 473) levels are lower after Torin1 and higher after rapamycin treatment.

Ganetespib at lower doses, up to 10 nM, had little or no effect on expression of key mTOR signaling proteins, or activation of mTORC1, as assessed by pS6 (Ser240/244) levels (**Fig 6**). At 100 nM and 1 µM, there was a major decrease in expression of mTOR, AKT, and pS6 (Ser240/244) expression in all cell lines (**Fig 6**).

**Fig 6.**
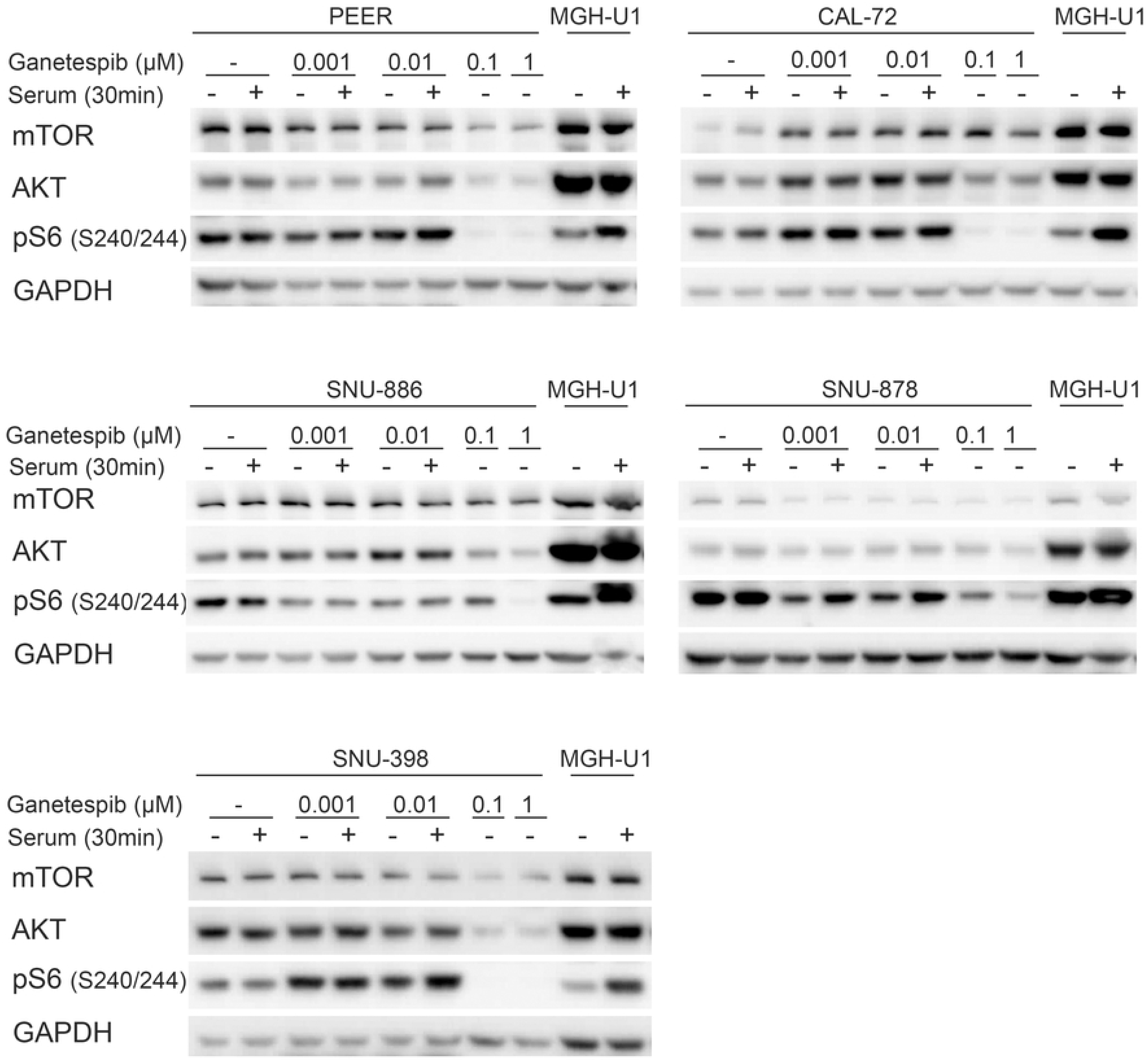
Effect of ganetespib on protein expression and mTORC1 signaling in TSC-mutant cell lines. Cells were treated with ganetespib in increasing concentrations (1, 10, 100 nM, and 1 µM) for 24h. Cells were serum starved for 24h (−) or after starvation had serum add back for 30 min (+). Effects on mTOR, AKT, and pS6 (Ser240/244) expression were seen at 0.1 and 1mM.

We also examined the effects of ganetespib on phosphorylation of 4E-BP1 and induction of apoptosis as assessed by cleaved caspase 3 levels (51) (**Fig 7**). Ganetespib at 100nM had little or no effect on phosphorylation of 4E-BP1; while rapamycin 20nM had variable effects in different cell lines, and Torin1 completely eliminated phosphorylation in all cell lines, as assessed by mobility shift (**Fig 7**). This lack of effect of ganetespib is consistent with previous studies (50). Cleaved caspase 3 levels varied widely among these 5 cell lines. PEER cells had high levels of cleaved caspase 3 under all conditions, likely due to growth in suspension culture, such that dead cells could not be eliminated. Nonetheless cleaved caspase 3 levels were increased at high doses of ganetespib in PEER cells. SNU-886 showed a major increase in cleaved caspase 3 levels after ganetespib treatment, and no or minimal effect from rapamycin or Torin1. SNU-878 cells showed a major increase in cleaved caspase 3 levels after each of rapamycin, Torin1, and ganetespib. CAL-72 and SNU-398 cells showed no increase in cleaved caspase 3 in response to any treatment.

**Fig 7.**
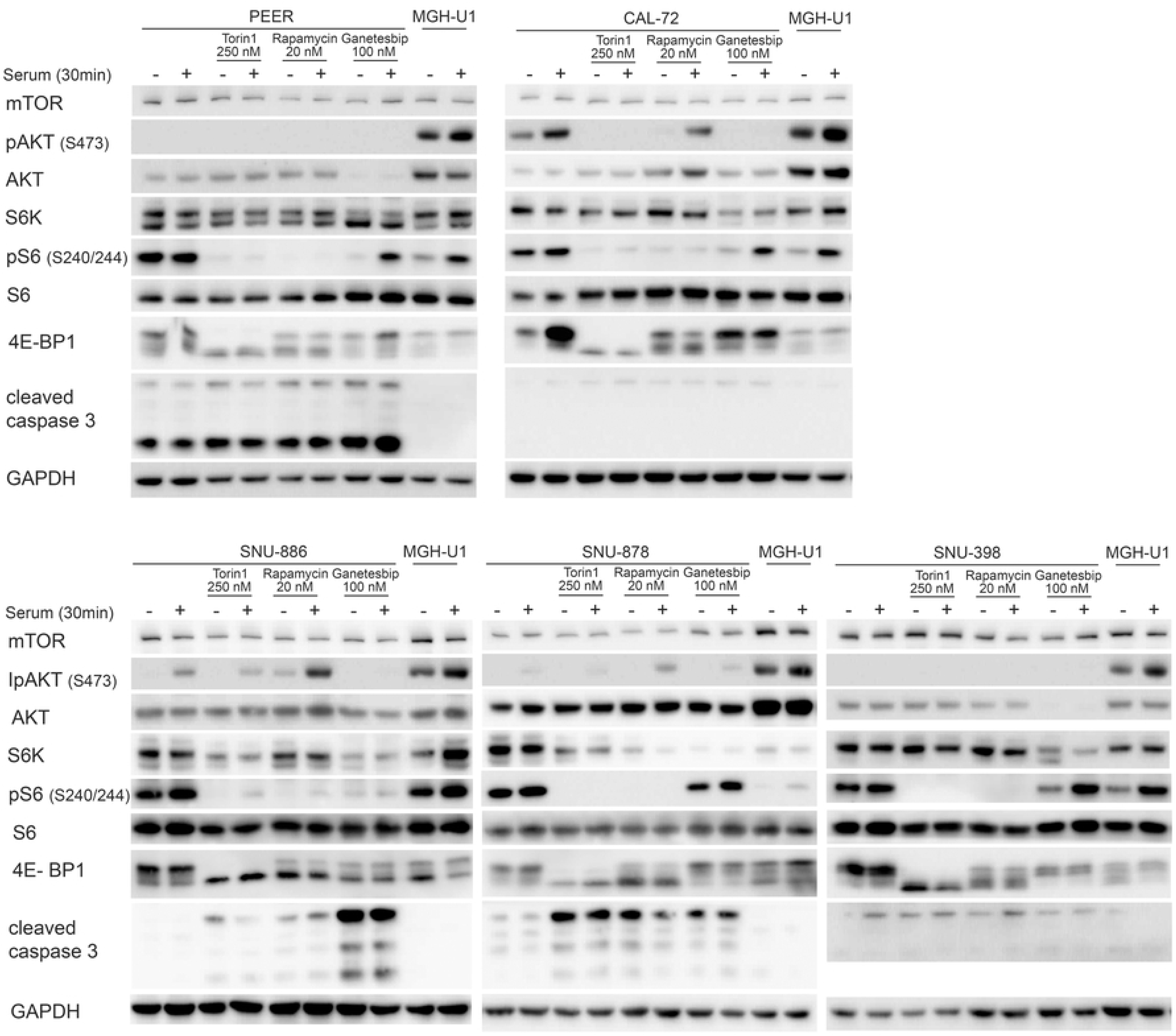
Impact of rapamycin, Torin1, and ganetespib on constitutively activated mTOR signaling and cleaved caspase 3. Cells were treated with rapamycin (20 nM), Torin1 (250 nM) or ganetespib (100 nM) for 24h. Cells were serum starved for 24h (−) or received after starvation serum add back for 30 min (+). Cells treated with rapamycin do not show expression of pS6 (Ser240/244) and expression of p4E-BP1 isoforms are reduced due to mTORC1 inhibition. Torin1-treated cells show no pS6 (Ser240/244), p4E-BP1, and a lowered pAKT (Ser473) expression, due to mTORC1/2 inhibition. Ganetespib-treated cells show a lowered AKT, S6K, and pS6 (Ser240/244) expression. The expression of cleaved caspase 3 is very distinctive among different cell lines.

The observation that there was little effect on mTORC1 signaling or protein expression in any of the cell lines near the IC50 dose of ganetespib (3 to 35nM) suggests that the growth inhibition effects were being mediated by other client proteins of HSP90 whose expression was likely reduced to some extent at doses near 10nM, and likely a collective effect on multiple HSP90 client proteins.

### Mouse xenograft tumor model studies

Given the evidence of some synergy in the growth inhibition of the TSC1/TSC2 null cell lines in response to combined mTOR and HSP90 inhibition, and evidence that they were impacting growth through different mechanisms, we explored the potential synergistic effect of treatment with these compounds in vivo using a subcutaneous xenograft tumor model with SNU-398 cells. 3.0×10^6^ SNU-398 cells were injected subcutaneously into the flank region of immunodeficient CB17SC-M (C.B-*Igh-1^b^*/IcrTac-*Prkdc^scid^*) mice. After approximately 10 days palpable and measurable tumors with a diameter of 3-5 mm were noted. Mice were treated with rapamycin, INK 128, or ganetespib, using doses described previously as being the maximal tolerated dose, in studies by ourselves and others (30, 48).

Mice were treated when tumors had a mean (SD) diameter of 7.8 (1.7) mm and a mean (SD) volume of 186 (110) mm^3^, after a mean (SD) of 18 (4.8) days after tumor cell injection.

Treatment with the individual drugs was assigned randomly.

Growth of xenograft tumor nodules was reduced by each of the three drugs, with ganetespib, 50 mg/kg by tail vein injection 1 time/week, having the least effect; and rapamycin, 3mg/kg given intra-peritoneally 3 days per week, having the greatest effect; and INK 128, 1 mg/kg by gavage 5 days/week, being intermediate (**Fig 8a and b, S22 a-d Fig**).

**Fig 8.**
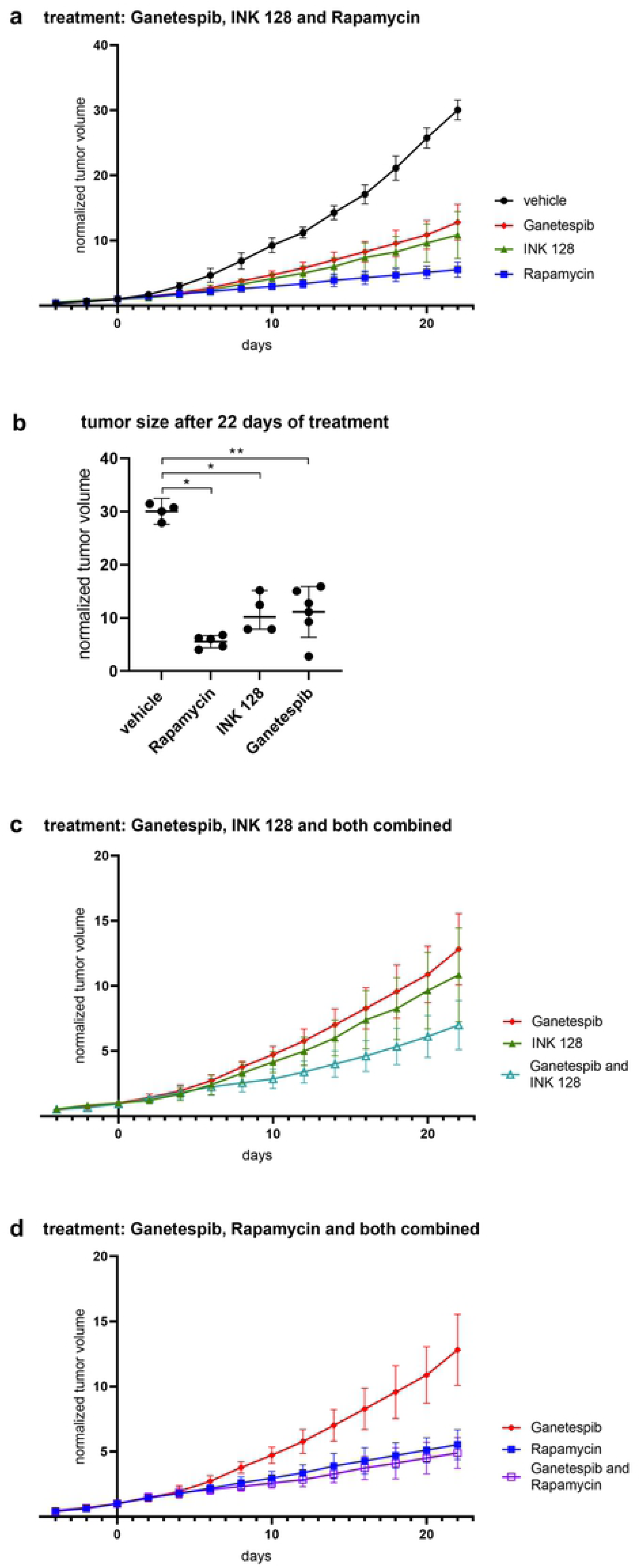
Xenograft tumor growth under vehicle, rapamycin, INK 128, ganetespib (a, b) combined ganetespib and INK 128 (c) and ganetespib and rapamycin (d) treatment. Tumor volume is shown as normalized to tumor volume at day 1 of treatment. Tumor size is shown as mean and standard deviation. Normalized tumor size of different treatment groups was compared on day 22 using Wilcoxon Rank Sum test. P-values less than 0.05 were considered statistically significant. Mice were treated with rapamycin (3 mg/kg, 3x/week, i.p., n=5), INK 128(1mg/kg, 5x/week, i.g., n=5) or ganetespib (50mg/kg, 1x/week, i.v., n=9) (a) or in combination of ganetespib and INK 128 (n=5) (c) or ganetespib and rapamycin (n=6) (d). Tumor size of treated mice is significantly smaller on day 22 compared to mice, which received vehicle (n=8) (b).

We then examined the potential benefit of combination treatment, with ganetespib and rapamycin, or with ganetespib and INK 128, in this xenograft model system. Combination ganetespib-rapamycin had similar effects on growth as rapamycin alone (**Fig 8d**). In contrast, ganetespib-INK 128 showed apparent synergy with a greater effect on growth than either drug alone although this did not achieve statistical significance (**Fig 8 c, S22e** and **f Fig**). None of the treated mice showed ill effects from treatment, including weight loss >10%, or skin lesions.

We also examined the effects of the drug treatments on mTOR signaling in vivo, through analysis of tumors by immunohistochemistry (IHC), and immunoblotting of tumor lysates. Histologically, the tumors were not homogeneous, with necrotic areas centrally and high proliferation in the periphery of nodules, regardless of treatment, as is commonly seen in xenograft models.

IHC against proliferating cell nuclear antigen (PCNA) was used to assess proliferation (52), and the ApopTag kit was used to detect apoptotic cells by the TUNEL method (53). IHC showed no TSC2 expression in the tumors. Vehicle and ganetespib-treated tumors were pS6 (Ser235/236)+. In contrast rapamycin-treated tumors had much reduced pS6 (Ser235/236) expression, and this was also lower in INK 128-treated tumors, in comparison to the tumors treated with ganetespib or vehicle (**Fig 9**).

**Fig 9.**
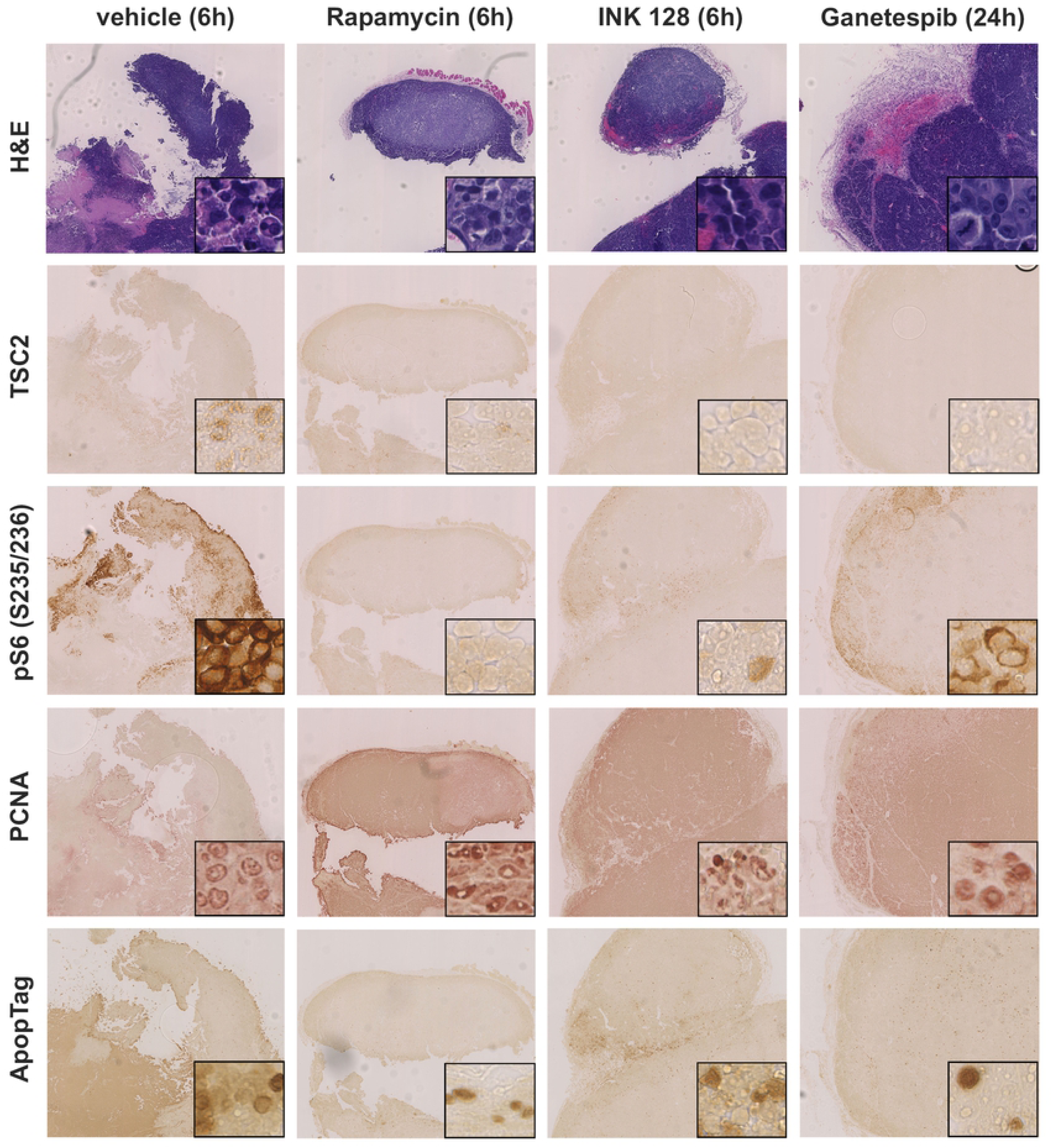
Immunohistochemical analysis of xenograft tumors of mice treated with vehicle, INK 128, ganetespib, or rapamycin for 21 days. Xenograft tumors generated from SNU-398 cells were harvest 24h after ganetespib treatment, 6h after INK 128 and rapamycin treatment, and stained using H&E, pS6 (S235/236), TSC2, ApopTag, or PCNA antibodies. Images shown were 60X magnified, insets showed portions of the tumor at higher magnification (400X). Tumors showed a distinct vascularization. TSC2 was not expressed in any tumor. PS6 (S235/236) expression was stronger in vehicle- and ganetespib-treated mice and correlated with locations of higher proliferation.

Immunoblots of harvested tumors, carefully dissected for viable and not necrotic regions, and livers, of mice treated with vehicle, rapamycin, INK 128, ganetespib, and the combinations were performed to examine the effect of the drugs on protein expression and the mTORC1 pathway in vivo. None of the treated livers showed major changes in expression of AKT or S6K (**Fig 10**). Similarly S6 and 4EB-P1 levels were similar under all treatments, with the exception of the ganetespib and INK 128 combination, which reduced both considerably (**Fig 10**). TSC2 expression was low in all xenografts, as expected, with some expression seen likely due to ingrowing vessels and connective tissue. pAKT (Ser473) expression was also universally low, as expected (**Fig 10**). Both rapamycin alone and in combination with ganetespib abolished pS6K (Thr389) and pS6 (Ser240/244) expression; while INK 128 alone or in combination reduced pS6 (Ser240/244) and eliminated pS6K (Thr389) (**Fig 10**). None of the treatments affect 4E-BP1 isoform expression, while the ganetespib and INK 128 combination reduced overall 4E-BP1 expression. Livers from the treated mice showed similar effects as the xenograft nodules, from each drug, except that pS6 (Ser240/244) was relatively highly expressed in the INK 128 treated liver (**Fig 10**).

**Fig 10.**
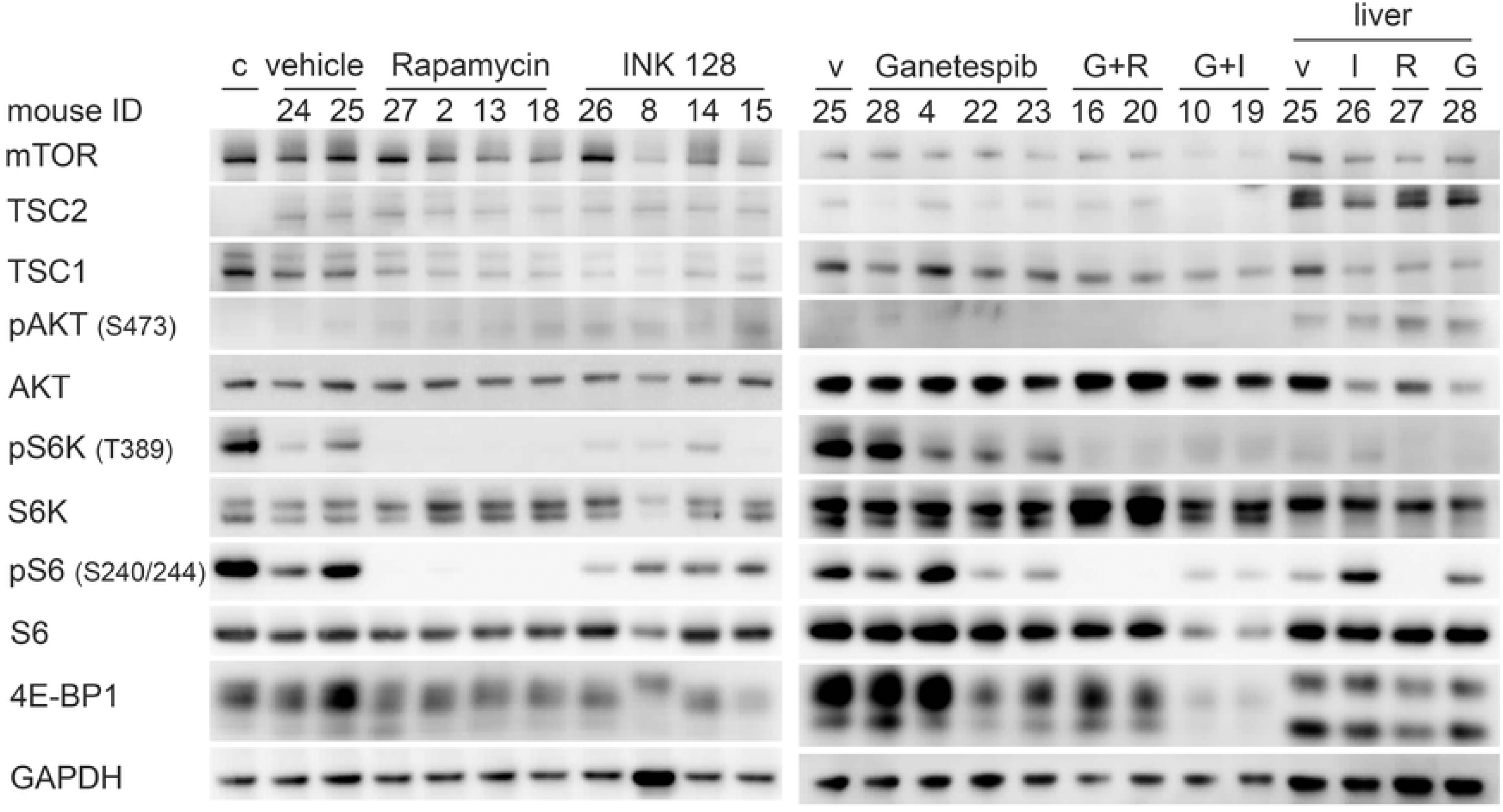
Effects of rapamycin, INK 128, ganetespib, and combination treatment on expression and mTOR signaling in SNU-398 xenograft tumor and liver cells. Mice were treated with rapamycin (3 mg/kg, 3x/week, i.p.), INK 128 (1mg/kg, 5x/week, p. o.), ganetespib (50mg/kg, 1x/week, i.v.) or in combination. Tumors were harvest and lysed 24h after ganetespib treatment and 6h after INK 128 or rapamycin treatment. Rapamycin treatment inhibited pS6K (Thr 389) and pS6 (Ser240/244) expression in the tumors. INK 128 treatment reduced pS6K (Thr 389), pS6 (Ser240/244), and p4E-BP1 isoform expression. Ganetespib reduced pS6K (Thr 389) and pS6 (Ser240/244) expression in some tumors. Combined ganetespib and rapamycin or INK 128 treatment showed a stronger inhibition of the mTOR pathway than each drug by itself. Rapamycin even inhibited pS6 (Ser240/244) expression in the liver. *Abbreviations: c: cells from cell culture, v: vehicle, G: Ganetespib, I: INK 128, R: Rapamycin*.

## Discussion

The current and accelerating trend in cancer therapy is the use of personalized (also called targeted or directed) cancer treatment. Such targeted therapies may be directed at genetic mutations which are driver events in cancer; or at expressed proteins, such as the estrogen receptor in breast cancer (54).

One resource that has been developed for a better understanding of cancer development, and to investigate potential targeted therapies are cancer cell lines, including the Cancer Cell Line Encyclopedia of 947 human cancer cell lines (55). In this study we searched for cancer cell lines with bi-allelic inactivating mutations in *TSC1* or *TSC2* from this resource. From a large set of candidate cell lines, subjected to sequencing and immunoblotting, we identified 5 cell lines with a total loss of either TSC1 or TSC2, and which also showed constitutively upregulated mTORC1 signaling, as indicated by high expression of pS6K (Thr389) and pS6 (Ser240/244) in the absence of serum. Two other cell lines, MFE-319 and OVK18, also showed constitutively upregulated mTORC1 signaling as well as upregulated pAKT (Ser473) expression, likely due to loss of PTEN by mutation (31, 32).

The mTORC1 signaling pathway is one of the main regulators of cell growth, which acts by enhancing anabolic biosynthetic pathways in both normal and cancer cells. The mTOR signaling pathway is involved in many important processes including proliferation, autophagy, protein and lipid synthesis (4, 56), lysosome and ribosome biogenesis, glucose and mitochondrial metabolism (57), and angiogenesis (58). As a key regulator of cell growth, the mTOR pathway is regulated by many upstream signals including hypoxia, inflammation, growth factors, DNA damage, energy deficiency, and nutrients (3). Both the PI3K and MAPK/Erk pathways influence activation of mTORC1, with signaling through PI3K and downstream elements having the predominant effect.

mTOR is an atypical serine/threonine kinase, and forms the key component of two protein complexes, mTORC1 and mTORC2. The TSC1 and TSC2 proteins function in a complex with a third component, TBC1D7 (9), as one major regulator of mTORC1. The TSC protein complex functions as a GTPase activating protein for the small GTP binding protein RHEB, a member of the extended RAS family of proteins. The TSC2 protein contains the GAP domain for RHEB and TSC1 has a stabilizing function for TSC2, such that both TSC1 and TSC2 are absolutely required for the activity of the TSC protein as a GAP for RHEB. RHEB-GTP binds to and activates mTORC1 at the lysosomal membrane. Hence, loss of the TSC protein complex, either TSC1 or TSC2, leads to high levels of RHEB-GTP, and constitutive activation of mTORC1 (59).

Inactivating mutations in either *TSC1* or *TSC2* in the germ line cause Tuberous Sclerosis Complex, an autosomal dominant neurocutaneous disorder, characterized by skin lesions, neuropsychiatric disorders and mainly benign tumors of the brain, kidney, lung, heart, and skin (2, 4). Mutation and loss of function of *TSC1* and/or *TSC2* have also been shown to occur consistently in a variety of sporadic cancers including bladder, kidney, pancreatic neuroendocrine tumors, and PEComa. More rarely *TSC1/TSC2* mutations have been identified in many other cancer types, though whether they are important driver events or passenger mutations is not clear in many instances.

Following our identification of five cell lines, PEER, CAL-72, SNU-886, SNU-878, and SNU-398, with bi-allelic mutation and loss of either *TSC1* or *TSC2,* we performed drug screen using 197 kinase inhibitors. The cell lines were sensitive to several mTOR inhibitors, but also HSP90 and cell cycle inhibitors. In addition, PEER cells were sensitive to Aurora kinase inhibitors.

We showed that the sensitivity of these cell lines to mTOR inhibitors like Torin1 and INK 128 correlated strongly with the deficiency of TSC2 and could be reversed by add back of TSC2 expression. However, we did not see this correlation with WYE-125132, AZD8055, and Torin2, which may reflect the activities of these compounds on kinases other than mTOR. Various rapalogs have been approved for treatment of various cancers, but complete responses are rare, and the PR rate is typically 5-10% (60-62). In addition, there are several case reports of exceptional response to rapalogs for several different types of cancer, some of which have been durable, going on for several years. These include PEComa (6), renal cell carcinoma (63), bladder cancer (64), and anaplastic thyroid cancer (63, 65). The reason why some patients with TSC1 or TSC2 mutant cancers have dramatic responses to rapalogs while the vast majority do not, is not understood.

One potential reason for the limited therapeutic benefit of rapalogs, is their lack of effect on many mTORC1 phosphorylation targets, including 4E-BP1. In addition, rapalog therapy induces many feedback loops leading to re-activation of PI3K and AKT signaling (66). Here we demonstrated that rapamycin was very effective at inhibiting classic mTORC1 downstream targets, including S6K and S6 phosphorylation, and showed minimal reduction in 4E-BP1 phosphorylation in these TSC mutant cancer cell lines. The mTORC2 pathway was not inhibited by rapamycin, and AKT phosphorylation was upregulated in some cell lines, likely due to feedback and counter-regulatory pathways (66). Rapamycin treatment induced very little apoptosis, but rather suppressed cell growth.

Potentially, newer ATP kinase pocket directed mTOR inhibitors could or should be more effective in controlling the growth of cells and tumors with TSC1/TSC2 inactivation. Torin1, for example, showed complete inhibition of phosphorylated 4E-BP1 and AKT. However, an apoptotic effect was still lacking. In addition, in our xenograft assay, rapamycin was more effective than the dual mTORC1/2 inhibitor INK 128 in inhibition of growth of the TSC2 null S-398 cell line. However, this may reflect the consequence of dual mTORC1 and mTORC2 inhibition by INK 128, leading to a lower tolerable dose, and less effective suppression of each of mTORC1 and mTORC2. Other off-target effects of this drug may also contribute to in vivo toxicity, limiting effective targeting of mTOR.

The HSP90 inhibitors ganetespib and NVP-AUY922 both showed strong inhibition of growth of all five TSC1/TSC2 null cell lines in vitro, with IC50’s in the range of 2 to 35 nM. Ganetespib was studied in greater detail. Ganetespib reduced mTOR, AKT, and S6K, as well as pS6, levels in all cell lines, but at 100 nM, a dose higher than the IC50 dose. These results suggest that ganetespib’s effect was not due directly to effects on mTOR signaling, but rather likely to broad effects in lowering expression of HSP90’s client proteins.

Based on these findings of growth inhibition by ganetespib in vitro, we performed tumor xenograft experiments using the TSC2 null SNU-398 cell line. This cell line showed robust growth in immunodeficient CB17SC-M scid mice as subcutaneous xenografts. All three of rapamycin, INK 128, and ganetespib, when given at previously determined doses near the maximally tolerated dose, showed a significant effect on the growth of the SNU-398 xenografts (**Fig 8**). Rapamycin was the most effective drug in these experiments, while INK 128 and ganetespib had less effects on growth and were similar to each other. Notably, all 3 drugs caused a reduction in growth rate, but none showed tumor regression.

Next, we examined the potential of combination therapy using ganetespib with each of the other two drugs. Ganetespib combined with INK 128 showed greater reduction in tumor growth than either drug alone, although this did not quite meet statistical significance (**Fig 8c**). In contrast, ganetespib combined with rapamycin showed nearly identical effects in tumor growth inhibition to rapamycin alone, indicating no in vivo synergy from the combination (**Fig 8d**).

Multiple past studies have examined the interaction between mTOR signaling and HSP90, and the potential benefit of HSP90 inhibition on growth of cell lines lacking *TSC1* or *TSC2.* Blenis and colleagues found that the combination of glutaminase (GLS) and HSP90 inhibition selectively triggered the death of TSC1/TSC2 deficient cells (67), likely due to the combination of oxidative and proteotoxic stress. Mollapour and colleagues reported that TSC1 was a co-chaperone of HSP90, facilitating HSP90 function to enhance proper folding of client proteins, including TSC2 (68). They then went on to show that loss of TSC1 leads to reduced acetylation of HSP90 at K407/K419, which leads to decreased binding by ganetespib; and that inhibition of histone deacetylases with concurrent ganetespib treatment led to enhanced growth suppression of RT4, a TSC1 null bladder cancer cell line (69).

In addition, the combination of rapamycin and HSP90 inhibitors has been previously studied for treatment of hepatocellular cancer (HCC) (69). Lang et al. reported that the combination of rapamycin and 17-(dimethylaminoethylamino)-17-demethoxygeledanamycin (17-DMAG), an HSP90 inhibitor, had a greater effect than either drug alone in reducing the growth rate of Huh-7 cells, an HCC cell line, in a subcutaneous xenograft model (69). However, the difference in comparison to single drug treatments was modest, and the combination reduced tumor growth without causing reduction in tumor size. They also studied a syngeneic orthotopic model of HCC, using a mouse HCC cell line, Hepa129. In that model, the rapamycin - 17-DMAG combination showed a dramatic synergistic effect in reducing tumor volume (69). There are many differences between this study and ours reported here, including different cell lines, different agents and doses being studied, and use of a syngeneic orthotopic model, that may explain these distinct results.

Numerous inhibitors of HSP90 have been developed as potential anticancer drugs. Many such drugs, including ganetespib, have been evaluated in clinical trials, but none have been approved by the FDA, due to lack of benefit. The main reason for the limited efficacy appears to be that at effective doses of HSP90 inhibition, there is release of the HSF1 transcription factor from HSP90, which enters the nucleus leading to a prosurvival heat shock response (70). Nonetheless clinical trials of HSP90 inhibitors often combined with other agents continue (see https://www.clinicaltrials.gov/).

In conclusion, we have identified and validated five cancer cell lines that have complete loss of either *TSC1* or *TSC2*, and consequent constitutive mTORC1 activation. Through our kinase inhibitor screen, we identified a number of inhibitors that have some activity on these cell lines, including cell cycle kinase and HSP90 kinase inhibitors. In vitro, the HSP90 inhibitors NVP-AUY922 and ganetespib had major inhibitory effects on the growth of all five lines in the nanomolar range. However, this appeared to be due to global effects on HSP90 client protein expression, and not a specific effect on components of the mTOR signaling pathway. In vivo analysis of an HCC subcutaneous xenograft model using the TSC2 null SNU-398 cell line showed that ganetespib at usual doses had minimal effects on tumor growth both alone and in combination with rapamycin and INK 128. Rapamycin showed activity superior to that of INK 128 in vivo.

## Acknowledgements

All cell lines were obtained from the Broad Institute. June Goto, Magdalena Tyburczy, Damir Khabibullin, and Yvonne Chekaluk provided assistance with procedures; Nathanel Gray provided the LINCS library.

## S Fig Legends

**S1 Fig. Confirmation of reported mutations by Sanger sequencing**

Sequencing traces with mutation are shown, including a control for each sequenced region. PEER cells showed a homozygous nonsense mutation in TSC1 and SNU-878 and SNU-886 cells a homozygous nonsense mutation in TSC2. All other cell lines showed heterozygous mutations.

**S2 Fig. IC50 determination for Rapamycin**

Rapamycin was serially diluted three-fold from 10 μM to 1.5 nM. Cell viability was determined 72h after treatment using CellTiter-Glo and XLfit4.0 software. Cell viability is shown relative to control, n= 2-4. Rapamycin did not achieve IC50 over this dose range.

**S3 Fig. Cell viability after WYE-125132 treatment**

WYE-125132 was serially diluted three-fold from 10 μM to 1.5 nM. Cell viability was determined 72h after treatment using CellTiter-Glo and XLfit4.0 software. Cell viability is shown in relative control activity, n= 2-4. IC_50_ was calculated as the drug concentration that reduced cell viability by 50% compared to untreated cells. WYE-125132 is an mTORC1/2 inhibitor. The SNU cell lines and CAL-72 were very sensitive to all tested mTORC1/2 inhibitors.

**S4 Fig. Cell viability after AZD8055 treatment**

AZD8055 was serially diluted three-fold from 10 μM to 1.5 nM. Cell viability was determined 72h after treatment using CellTiter-Glo and XLfit4.0 software. Cell viability is shown in relative control activity, n= 2-4. IC_50_ was calculated as the drug concentration that reduced cell viability by 50% compared to untreated cells. AZD8055 is an mTORC1/2 inhibitor. The SNU cell lines and CAL-72 were very sensitive to all tested mTORC1/2 inhibitors.

**S5 Fig. Cell viability after Torin1 treatment**

Torin1 was serially diluted three-fold from 10 μM to 1.5 nM. Cell viability was determined 72h after treatment using CellTiter-Glo and XLfit4.0 software. Cell viability is shown in relative control activity, n= 2-6. IC_50_ was calculated as the drug concentration that reduced cell viability by 50% compared to untreated cells. Torin1 is an mTORC1/2 inhibitor. The SNU cell lines and CAL-72 were very sensitive to all tested mTORC1/2 inhibitors.

**S6 Fig. Cell viability after Torin2 treatment**

Torin2 was serially diluted three-fold from 10 μM to 1.5 nM. Cell viability was determined 72h after treatment using CellTiter-Glo and XLfit4.0 software. Cell viability is shown in relative control activity, n= 2-4. IC_50_ was calculated as the drug concentration that reduced cell viability by 50% compared to untreated cells. Torin2 is an mTORC1/2 inhibitor. Among the tested mTOR inhibitors, Torin2 showed the lowest IC50 for each of the 5 cell lines.

**S7 Fig. Cell viability after INK 128 treatment**

INK 128 was serially diluted three-fold from 10 μM to 1.5 nM. Cell viability was determined 72h after treatment using CellTiter-Glo and XLfit4.0 software. Cell viability is shown in relative control activity, n= 2-4. IC_50_ was calculated as the drug concentration that reduced cell viability by 50% compared to untreated cells. INK 128 is an mTORC1/2 inhibitor. All TSC1 or TSC2 deficient tumor cell lines were very sensitive to all tested mTORC1/2 inhibitors.

**S8 Fig. Cell viability after Flavopiridol treatment**

Flavopiridol was serially diluted three-fold from 10 μM to 1.5 nM. Cell viability was determined 72h after treatment using CellTiter-Glo and XLfit4.0 software. Cell viability is shown in relative control activity, n= 2-4. IC_50_ was calculated as the drug concentration that reduced cell viability by 50% compared to untreated cells. Flavopiridol is a CDKs inhibitor. All cell lines were sensitive to Flavopiridol.

**S9 Fig. Cell viability after CGP60474 treatment**

CGP60474 was serially diluted three-fold from 10 μM to 1.5 nM. Cell viability was determined 72h after treatment using CellTiter-Glo and XLfit4.0 software. Cell viability is shown in relative control activity, n= 2-4. IC_50_ was calculated as the drug concentration that reduced cell viability by 50% compared to untreated cells. CGP60474 is CDKs and mTOR inhibitor. All cell lines were sensitive to CGP60474.

**S10 Fig. Cell viability after BMS-387032 treatment**

BMS-387032 was serially diluted three-fold from 10 μM to 1.5 nM. Cell viability was determined 72h after treatment using CellTiter-Glo and XLfit4.0 software. Cell viability is shown in relative control activity, n= 2-4. IC_50_ was calculated as the drug concentration that reduced cell viability by 50% compared to untreated cells. BMS-387032 is a CDKs inhibitor. SNU-398 cells were the most sensitive cell line to BMS-387032.

**S11 Fig. Cell viability after GSK461364 treatment**

GSK461364 was serially diluted three-fold from 10 μM to 1.5 nM. Cell viability was determined 72h after treatment using CellTiter-Glo and XLfit4.0 software. Cell viability is shown in relative control activity, n= 2-4. IC_50_ was calculated as the drug concentration that reduced cell viability by 50% compared to untreated cells. GSK461364 is a PLK inhibitor. The cell lines SNU-886, SNU-878, and SNU-398 were sensitive to GSK461364, while in contrast the cell lines PEER and CAL-72 were not sensitive.

**S12 Fig. Cell viability after HMN-214 treatment**

HMN-214 was serially diluted three-fold from 10 μM to 1.5 nM. Cell viability was determined 72h after treatment using CellTiter-Glo and XLfit4.0 software. Cell viability is shown in relative control activity, n= 2-4. IC_50_ was calculated as the drug concentration that reduced cell viability by 50% compared to untreated cells. HMN-214 is a PLK inhibitor. PEER cells were most sensitive to HMN-214.

**S13 Fig. Cell viability after GSK1070916 treatment**

GSK1070916 was serially diluted three-fold from 10 μM to 1.5 nM. Cell viability was determined 72h after treatment using CellTiter-Glo and XLfit4.0 software. Cell viability is shown in relative control activity, n= 2. IC_50_ was calculated as the drug concentration that reduced cell viability by 50% compared to untreated cells. GSK1070916 is an Aurora A, B and C inhibitor. TSC1 null PEER cells were very sensitive to all Aurora inhibitors, in contrast to TSC2 null SNU-398 cells, which were much less sensitive to Aurora inhibitors.

**S14 Fig. Cell viability after ZM-447439 treatment**

ZM-447439 was serially diluted three-fold from 10 μM to 1.5 nM. Cell viability was determined 72h after treatment using CellTiter-Glo and XLfit4.0 software. Cell viability is shown in relative control activity, n= 2. IC_50_ was calculated as the drug concentration that reduced cell viability by 50% compared to untreated cells. ZM-447439 is an Aurora A and B inhibitor. TSC1 null PEER cells were very sensitive to all Aurora inhibitors, in contrast to TSC2 null SNU-398 cells, which were much less sensitive to Aurora inhibitors.

**S15 Fig. Cell viability after AZD1152-HQPA treatment**

AZD1152-HQPA was serially diluted three-fold from 10 μM to 1.5 nM. Cell viability was determined 72h after treatment using CellTiter-Glo and XLfit4.0 software. Cell viability is shown in relative control activity, n= 2. IC_50_ was calculated as the drug concentration that reduced cell viability by 50% compared to untreated cells. AZD1152-HQPA is an Aurora A, B and C inhibitor. TSC1 null PEER cells were very sensitive to all Aurora inhibitors, in contrast to TSC2 null SNU-398 cells, which were much less sensitive to Aurora inhibitors.

**S16 Fig. Cell viability after XMD16-144 treatment**

XMD16-144 was serially diluted three-fold from 10 μM to 1.5 nM. Cell viability was determined 72h after treatment using CellTiter-Glo and XLfit4.0 software. Cell viability is shown in relative control activity, n= 2. IC_50_ was calculated as the drug concentration that reduced cell viability by 50% compared to untreated cells. XMD16-144 is an Aurora A and B inhibitor. TSC1 null PEER cells were very sensitive to all Aurora inhibitors, in contrast to TSC2 null SNU-398 cells, which were much less sensitive to Aurora inhibitors.

**S17 Fig. Cell viability after NVP-AUY922 treatment**

NVP-AUY922 was serially diluted three-fold from 10 μM to 1.5 nM. Cell viability was determined 72h after treatment using CellTiter-Glo and XLfit4.0 software. Cell viability is shown in relative control activity, n= 2-4. IC_50_ was calculated as the drug concentration that reduced cell viability by 50% compared to untreated cells. NVP-AUY922 is an HSP90 inhibitor. All cell lines were very sensitive to NVP-AUY922.

**S18 Fig. Cell viability after ganetespib treatment**

Ganetespib was serially diluted three-fold from 10 μM to 1.5 nM. Cell viability was determined 72h after treatment using CellTiter-Glo and XLfit4.0 software. Cell viability is shown in relative control activity, n= 2-4. IC_50_ was calculated as the drug concentration that reduced cell viability by 50% compared to untreated cells. Ganetespib is an HSP90 inhibitor. All cell lines were very sensitive to ganetespib.

**S19 Fig. Cell viability after GSK2126458 treatment**

GSK2126458 was serially diluted three-fold from 10 μM to 1.5 nM. Cell viability was determined 72h after treatment using CellTiter-Glo and XLfit4.0 software. Cell viability is shown in relative control activity, n= 2. IC_50_ was calculated as the drug concentration that reduced cell viability by 50% compared to untreated cells. GSK2126458 is a PI3K inhibitor. CAL-72 und the SNU cell lines were sensitive to GSK2126458.

**S20 Fig. Cell viability after WZ3105 treatment**

WZ3105 was serially diluted three-fold from 10 μM to 1.5 nM. Cell viability was determined 72h after treatment using CellTiter-Glo and XLfit4.0 software. Cell viability is shown in relative control activity, n= 2. IC_50_ was calculated as the drug concentration that reduced cell viability by 50% compared to untreated cells. WZ3105 is a CLK2, CNSK1E, FLT3 and ULK1 inhibitor. PEER, CAL-72 and SNU-398 cells were sensitive to WZ3105.

**S21 Fig. Cell viability after MK 1775 treatment**

MK 1775 was serially diluted three-fold from 10 μM to 1.5 nM. Cell viability was determined 72h after treatment using CellTiter-Glo and XLfit4.0 software. Cell viability is shown in relative control activity, n= 2-5. IC_50_ was calculated as the drug concentration that reduced cell viability by 50% compared to untreated cells. MK 1775 is a Wee1 inhibitor. All cell lines were sensitive to MK 1775.

**S22 Fig. S-398 tumor xenograft treatment.**

Tumor volume is shown as normalized tumor volume to day 1 of treatment. Tumors were measured every 2-3 days. Each tumor is depicted separately, n= 5-9 per treatment. Mice were treated with vehicle (a), ganetespib (50mg/kg, 1x/week, i.v.) (b), INK 128 (1mg/kg, 5x/week, i.g.) (c), rapamycin (3 mg/kg, 3x/week, i.p.) (d), or ganetespib and rapamycin combined (same doses) (e) or ganetespib and INK 128 combined (same doses) (f). Tumors under treatment grew less compared to vehicle-treated tumors.

